# A Dose-Response Model for Accurate Detection and Quantification of Transcriptome-Wide Gene Knockdown for Oligonucleotide-Based Medicines

**DOI:** 10.1101/2024.05.28.596270

**Authors:** David Pekker, Steven Kuntz, Monica McArthur, Tim Nicholson-Shaw, Sara Yanke, Swagatam Mukhopadhyay

## Abstract

Synthetic antisense oligonucleotides and siRNAs are a class of Oligonucleotide-Based Medicines (OBMs) that can hybridize with pre-mRNA and mRNA, recruit a mechanism-of-action specific enzymatic complex, and knockdown target gene expression. This class of molecules provides an excellent substrate for designing precision gene-modulatory therapeutics; however, quantifying on- and off-target dose response as measured by next-generation sequencing for this class of therapeutics has remained under-powered and ambiguous. Often *in silico* predictions of off-targets (ranked by edit tolerance) are used as putative off-target analysis in ASO and siRNA drug design. We construct a simple, effective theory of transcriptional dynamics and enzymatic activity in order to describe the transcriptome-wide response to these oligonucleotides. We establish rigorous quantification methods of off-target analysis in oligonucleotide drug design. We also extend the DESeq work [1, 2] of Negative Binomial noise in gene expression measurements to describe noise, including outliers, in OBM-dose response NGS experiments. We demonstrate the performance of our model on both synthetic and experimental Digital Gene Expression (DGE) data of dose response in ASO-treated cells. We present our analysis package, *DoReSeq*, as a freely available resource for the community. We hope this will elevate the standards of off-target analysis for such an important class of precision therapeutics.

## I. INTRODUCTION

Synthetic Antisense Oligonucleotides (ASOs) and small interfering RNAs (siRNAs) have ushered in a new era of precision medicine, offering promising avenues for the development of gene-specific therapeutics [3]. These molecules, designed to hybridize specifically with pre-messenger RNA (pre-mRNA) and messenger RNA (mRNA), exploit endogenous cellular mechanisms to modulate gene expression. By employing the right chemical composition, these synthetic oligonucleotides are able to recruit an enzymatic complex (for example, RNase H1) that initiates the knockdown of gene expression through an enzymatic cleavage reaction. This capability not only highlights their potential as powerful tools for precise gene-expression modulation but also underscores their significance in therapeutic applications [4]. ASOs and siRNAs are two important clinically-proven therapeutic modalities in the broader class of Oligonucleotide-Based Medicines (OBMs) [4].

The advantages of these gene-specific therapeutics, however, is contingent upon a nuanced measurement and quantification of their on-target and off-target effects, calling for a comprehensive approach to study their doseresponse relationships. Off-targets are unintended gene knockdowns or up-regulation owing to OBM treatment. With falling costs of Next-Generation Sequencing (NGS) methods, transcriptome-wide measurements, for example, Digital Gene Expression (DGE, see [5]) has become quite cost-effective for routine pre-clinical lead identification experiments, such as a well-sampled dose-response experiment in cell lines. Traditionally, dose-response has been quantified using gene-specific qPCR experiments where probes were designed for putative (*in silico* predicted) off-targets, limiting the scope of discovery of the true off-target liability of these lead molecules. These *in silico* methods rely on hybridization-dependent off-target assessments using genomics tools, tolerating up to one or two edits across the transcriptome [6]. However, this analysis has a crucial flaw: the true off-targets observed for ASOs and siRNAs are often not captured by *in silico* methods [7]. As a result, missed off-target effects remain a liability in developing siRNA and ASO drugs [8–10] and dampen the original hope of such precision medicine modalities as an “informational drug” [11].

In addition, dose-response curves for this class of molecules are usually fitted with the Hill equation. This approach is not obviously appropriate for oligo-mediated gene modulation. The Hill equation was motivated by receptor-ligand interactions and co-operativity of such interactions [12]. It is not *a priori* obvious that such a theory holds for ASO or siRNA induced enzymatic activity—for example, cooperative interaction in such reaction kinetics is debatable, and usually motivated by phenomenological modeling [13– Moreover, the widely accepted paradigm that frames gene knockdown detection as a least-square regression problem is not robust because gene expression noise is dependent on expression level. Such conventional regression methods restrict noise models to normal distribution regime— inapplicable in the context of Next-Generation Sequencing (NGS)-measured gene expression counts which from both biological and technical reasons appear to be Negative Binomial distributed [16– DESeq and DESeq2 introduced a much more nuanced approach to describing noise in gene expression counts [1, 2]. Specifically, Negative Binomial distributions were used to describe noise in counting measurements and a Bayesian modeling framework was used to robustly fit parameters using Maximum *A Posteriori* (MAP) estimation. This Bayesian framework was designed to adaptively manage the inherent complexities and variability of gene expression data, particularly those acquired through RNA sequencing (RNAseq) technologies, including 3’-quant-Seq variants [5], collectively referred to as Digital Gene Expression (DGE) from hereon [19].

In this paper, we describe DoReSeq (pronounced Dory-Seek), which stands for Dose-Response Model using Sequencing. DoReSeq is a method for characterizing the level of gene expression dependent upon the concentration of an ASO or an siRNA and as detected by NGS at every dose-point. Our goal is to both identify which genes are knocked down (whether on-target or off-target), as well as to quantify both the level of knockdown and our confidence in such knockdown calls.

DoReSeq is composed of three parts: a kinetic model of dose-response, a noise model that describes biological and technical noise in next generation sequencing assays, and a Bayesian inference fitter.

We first construct a kinetic, dose-responsive model of mRNA knockdown by an ASO. Our model describes how the competition between pre-mRNA maturation and the ASO guided cleavage of the pre-mRNA leads to a dosedependent time-evolution of the concentration of mature mRNAs. In the long-time limit our model predicts that the concentration of mRNAs as a function of the dose reaches a steady-state that coincides with the Monod equation (also known the empirical Hill equation with *n* = 1) [12]. Time between dosing and readout is an important confounding variable, that is often chosen to be different in different experimental setups. Therefore we intrinsically incorporate time into our dose-response model, which allows DoReSeq to be used to fit the RNA decay rate and thus compare results of experiments with different readout times.

Second, we supplement our kinetic model with a noise model of gene expression as measured by NGS. The noise model of DoReSeq is similar to that of DESeq/DESeq2 [1, 2], but applied to the realm of nonlinear models as opposed to generalized linear models. Specifically, we use our kinetic model to parameterize the mean response of a gene *i* as a function of dose *A* (and time *t*). We model noise using the Negative Binomial distribution that has a gene-specific dispersion parameter *ϕ*_*i*_ and a mean parameter, given by our kinetic model and scaled by a sample-determined scaling factor *s*_*α*_ (equal to the total number of non-duplicate reads for the sample, where *α* is the sample index).

Third, we use the full power of Bayesian inference to detect dose responsive genes and quantify their response from DGE datasets. Previous work on the use of Bayesian inference for analyzing gene expression focused on obtaining maximum *a posteriori* (MAP) estimates of model parameters [1, 2]. However, the recent advances in Markov Chain Monte Carlo (MCMC) sampling strategies [20], as well as increasing compute power, allow us to efficiently sample probability distributions of the DoReSeq model parameters across thousands of genes. As a result, DoReSeq is able to obtain not just the MAP estimates, but also credible regions and p-values for the model parameters and other key observables. We treat the sample-specific parameters: the scaling factor *s*_*α*_, the time *t*_*α*_, and the dose *A*_*α*_ as confounding/independent variables and *x*_*α,i*_ the raw number of reads of gene *i* in sample *α* as the dependent variable. We use MCMC to sample the DoReSeq model parameters for each gene *i*: (1) TPM_*i*_, the number of Transcripts per Million (TPM) at zero dose; (2) *ϕ*_*i*_, the negative binomial distribution dispersion parameter; (3) mKD_*i*_, the maximum knockdown at high dose; (4) rIC_50,*i*_, the relative rIC_50_, i.e. the dose needed to achieve a relative decrease in expression, as compared to mKD_*i*_, of 50%; and (5) *δ*_*i*_,the mRNA decay rate.

Using synthetic data, we show that we can use DoReSeq to robustly identify dose-responsive genes and recover the model parameters used to synthesize the data. By analyzing synthetic data, we discover an interesting caveat: for genes with weak dose-response, the mKD_*i*_ and rIC_50_ space contains one *stiff* and one *sloppy* direction [21]. The *sloppy* direction makes it hard to simultaneously fix the value of both mKD_*i*_ and rIC_50_. In Bayesian inference, this property results in credible regions being inflated along the *sloppy* direction. Somewhat counter-intuitively, we find that the knockdown at a specific dose (within the range of doses probed in the experiment) is a *stiff* observable. Therefore, in this regime DoReSeq is still able to robustly identify genes with weak dose-response, even though characterizing their response in terms of mKD_*i*_ and rIC_50_ is poor without additional data to pin down the *sloppy* direction.

Finally, we apply DoReSeq to experimental DGE data. As our dataset was acquired at a single time point, we were not able to fit *δ*_*i*_. Using DoReSeq, we were able to identify 203 dose-responsive genes, containing one ontarget response and 202 off-target responses (out of 8005 genes with good sequencing data) and obtain tight estimates of the model parameters for those genes.

By combining Negative Binomial distribution of gene expression noise with our dose-response model and Bayesian inference, our data fitting method accurately maps gene expression data to a family of most probable dose-response curves.

DoReSeq’s strength lies in its capacity to integrate large collection of DGE datasets pertaining to dose response analysis, especially beneficial in scenarios where a substantial number of control samples are available. It significantly increasing our confidence in the design and discovery of precision OBMs by providing a statistically sound method of identifying off-target liability. We make our software package freely available to the research community under MIT License [22], with our intention of promoting the application of these insights to the pre-clinical and clinical development in OBMs.

## II. THE DORESEQ KINETIC MODEL OF DOSE RESPONSE

In this section we construct our model that describes the competition between pre-mRNA maturation and ASO guided cleavage of the pre-mRNA on recruiting RNaseH. We focus on a single gene and consider the population of three species, see Fig. 1. We denote the population of pre-mRNAs by *T*, the population of premRNAs bound to an ASO by *T*^*∗*^, and the population of mature mRNAs by *M*. In our model, the ensemble of cells transcribes pre-mRNAs at a fixed, mean rate *β*. The pre-mRNAs can either mature, which occurs with rate *Tγ*, or become bound by an ASO, which occurs at the rate *TαA* where *A* is the ASO concentration and *α* is the rate constant. Once an ASO-pre-mRNA complex is formed it can either be cleaved, which occurs at rate *T*^*∗*^*κ*, or become a mature mRNA which occurs at rate *T*^*∗*^*γ*. Finally, the mature RNAs decay at rate *Mδ*. In constructing our kinetic model we are making the simplifying assumption that the ASO modulates gene expression exclusively through RNaseH mediated knockdown and that the ASO has no effect on the transcription rate *β*, the maturation rate *γ*, nor the mature RNA decay rate *δ*. As a consequence the description of biological processes such as ASO induced gene up-regulation is beyond the scope of our kinetic model. We also assume that ASOs are primarily active in the nucleus—a defensible assumption for intron-targeting ASOs and exon-targeting ASOs that are primarily active in the nucleus [23]. Our kinetic model has several key features, which we now discuss.

**FIG. 1.**
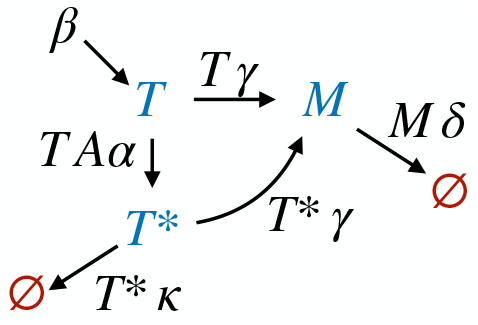
DoReSeq kinetic model diagram. The model describes population of three states *T, M*, and *T*^*∗*^ which corresponds to pre-mRNA, mature mRNA, and ASO bound premRNA. Arrows indicate transitions between states and labels over the arrows indicate the transition rates. Arrows pointing to *∅* indicate the destruction of the RNA. The arrow starting at *β* indicates the creation of mRNA at the mean transcription rate *β*.

One of the key results of our model is that knockdown can saturate to a finite, non-zero value, even at very high ASO concentrations. This feature emerges naturally because the model captures two critical biological mechanisms, in the tradition of simple systems-level models of biological processes [24]. First, we model a mechanism where the target-site may not be available for productive cleavage that leads to gene knockdown— because most ASOs act co-transcriptionally, this could be because of several RNA regulatory processes, for example, splicing removing the target-site into intronic lariats, RNA secondary structure, RNA-binding protein occupancy (RBPs), regulatory RNA occupancy reducing accessibility to target-site, etc. and therefore, to ASO-guided cleavage. Second, we made a simplifying assumption that the maturation rate of state *T* (regular pre-mRNAs) and state *T*^*∗*^ (ASO-bound pre-mRNAs) is identical. This assumption is reasonable because the ASO:RNA bound state is a very local stretch (10-20 Bp long) of the pre-mRNA, and the fate of the pre-mRNA without RNaseH activity is typically determined at a significantly longer length scale. Consequently, even at high ASO concentrations, where almost all pre-mRNAs become bound by ASOs, there is still a competition between the maturation and RNaseH-mediated-cleavage of bound pre-mRNAs. The fraction of bound pre-mRNAs that can be cleaved before they become mature sets the limit on the maximum knockdown.

In constructing our model, we have made the simplification that RNAs are transcribed at a steady rate. While this assumption is incorrect at a single-cell resolution for which transcription is known to occur in bursts [16], this a reasonable assumption for a large population of cells for which such transcriptional bursts average to an effective birth rate of pre-mRNA. We have made the simplification that that the off-rate of ASO is negligible compared to all other rates, meaning, there is no *T*^*∗*^ → *T* transition in our model. This assumption is justified for bridged-nucleic acid ASOs for which the melting temperature (of 16-20 mers) is rather high (for standard gapmer designs of phosphorothioate using Locked Nucleic Acids (LNA) [25]). It is easy to extend our model to include off-rate of ASO. Finally, we comment that in constructing our model we have explicitly ignored any potential mechanisms of cell perturbation induced by ASO treatment itself with broad effects on gene expression owing to, say, an apoptotic response. For example, cellular toxicity of ASOs can lead to up-regulation of the target gene in a regulatory-feedback pathway of cellular response. In DGE experiments, timescale of readout is important— such feedback typically happen at a longer timescale compared to enzymatic activity [26]. Our model includes kinetic effects to accommodate this biological challenge— the RNaseH-mechanism driven direct knockdown effects are easier to separate compared to indirect effects from ASO toxicity at earlier time-points (say, 5-6 hrs). However, at this time-point one may not reasonably expect for RNA knockdown effects to have reached steady state values because cellular uptake of ASOs and RNA halflife timescales are also in hours [27]. We see the kinetic (time-dependent) dose-response model as a major advantage of our work. To summarize, our model only includes first-order interactions for RNaseH-mediated knockdown.

We now write down the kinetic equations of our model and comment on their properties. The processes described above result in the following kinetic equations—

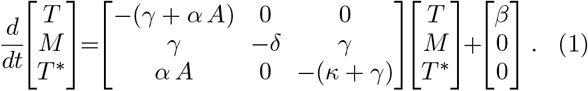

To analyze the implications of these rate equations we first find the steady-state populations in the absence of ASO treatment and then we use these steady-state populations as the initial condition for time evolution following ASO treatment.

To find the initial, steady-state populations *T*_0_, *M*_0_, and 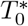, we set the LHS of eq. (1) to zero and also set the ASO concentration *A* = 0 to obtain

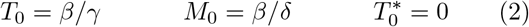

Using Eq. 2 as the initial conditions on Eq. (1), we find that the following closed form solution for population dynamics

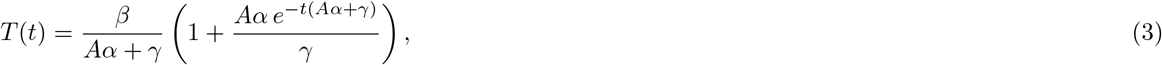

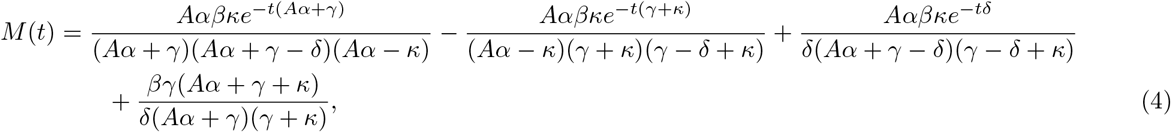

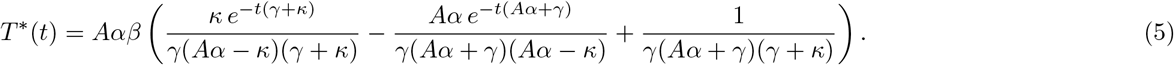

We now focus on the population of mature mRNAs, as that is the quantity that is being measured experimentally. Taking the long-time limit, we find the steady state population

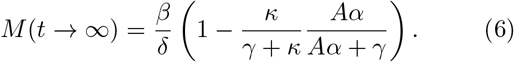

From this expression, we can read off the maximum knockdown (saturation effect) at high ASO concentration to be

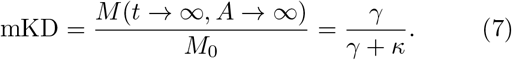

This expression highlights that the maximum knockdown is controlled by the competition of mRNA maturation rate *γ* and the ASO guided cleavage rate *κ*. For fitting experimental observations, we find it convenient to reparameterize the response using mKD and rIC_50_ = *γ/α*

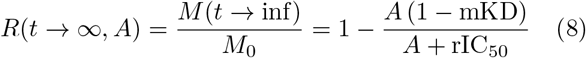

Let us now explore how *M* (*t*) approaches the steady state. The expression for the time dependence of *M* (*t*) in equation Eq. (5) is rather complicated, involving four different terms, one static and three exponential decays controlled by three different rates: *Aα* + *γ, γ* + *κ*, and *δ*. To gain intuition, we first look at the constituent rates *Aα, γ, κ*, and *δ*. In order for knockdown to occur *κ, Aα* ≳ *γ*, otherwise there will not be appreciable flow *T* → *T*^*∗*^ → cleavage. On the other hand we expect to be in the regime *δ* ≪ *γ*), which is usually appropriate for genes in eukaryotic cells in which mRNA maturation time is much shorter than mRNA half-life [28]. Hence we expect there to be a short transient associated with the fast rates *Aα* + *γ* and *γ* + *κ* in which the populations *T* (*t*) and *T*^*∗*^(*t*) reach their steady-state values, followed by a long decay associated with rate *δ* over which *M* (*t*) reaches its steady-state value. Which is indeed what we observe, see Fig. 2 a for the case *δ* = 0.2*γ*.

**FIG. 2.**
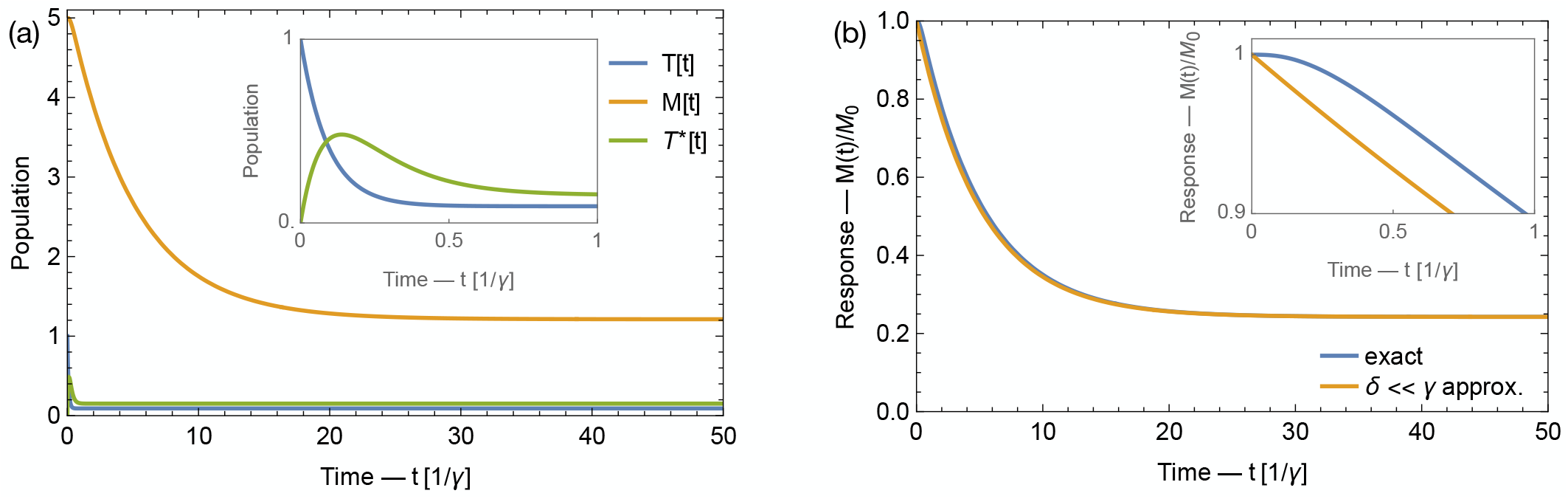
DoReSeq dynamics in the regime that mRNA decay rate is much smaller than mRNA maturation rate: *δ ≪ γ*. (a) Populations dynamics of pre-mRNA (*T* (*t*)), mature mRNA (*M* (*t*)), and ASO bound pre-mRNA (*T*^*∗*^(*t*)) as a function of time following ASO injection. Populations are in arbitrary units, time is in units of mRNA maturation rate *γ*. Inset shows a zoom in on populations *T* (*t*) and *T*^*∗*^(*t*) at short times. Observe that following the introduction of the ASO, the populations of *T* (*t*) and *T*^*∗*^(*t*) states quickly reach their steady-state values while the population of *M* (*t*) state takes much longer time to reach its steady-state value. (b) Mature mRNA response curves obtained with full dynamics and with the approximate *δ ≪ γ* dynamics. Inset shows zoom in on the initial dynamics. Observe that the two models predict similar dynamics except for a short-time transient which is absent in the approximate dynamics. [to produce this plot we used *α/γ* = 1, *A* = 5, *β/γ* = 1, *δ/γ* = 0.2, *κ/γ* = 5]

Next, we make the simplifying assumption that *δ* ≪ *γ*, i.e. we choose to ignore the initial transient in which the populations *T* (*t*) and *T*^*∗*^(*t*) reach their steady-state values. Under this assumption, we can drop the two fast decaying terms and simplify the denominator of the remaining term of *M* (*t*) in Eq. (5) to obtain the response function

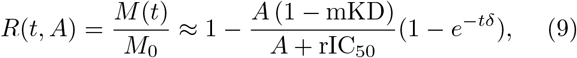

where we have used the same parametrization in terms of mKD and rIC_50_ as in Eq. (8). In Fig. 2 we compare the response function obtained from the exact time evolution Eq. (5) and from the approximate time evolution Eq. (9). We observe that the two curves are almost identical, except for the small initial transient that can be seen in the zoomed view plotted in the inset.

In Fig. 3 we plot response curves as a function of dose for various times using both the exact solution of Eq. (5) as well as the approximate solution Eq. (9). We observe that if we fix time, the response increases with dose saturating at high dose. At the same time we observe that as time increases so does the response until the steady state is reached at long times. Finally we observe that for *δ* ≲ 0.2*γ*, exact and approximate solutions of the kinetic model are almost identical. Eq. (9) is a simple and convenient expression for parameterizing the response of gene expression to ASO treatment.

**FIG. 3.**
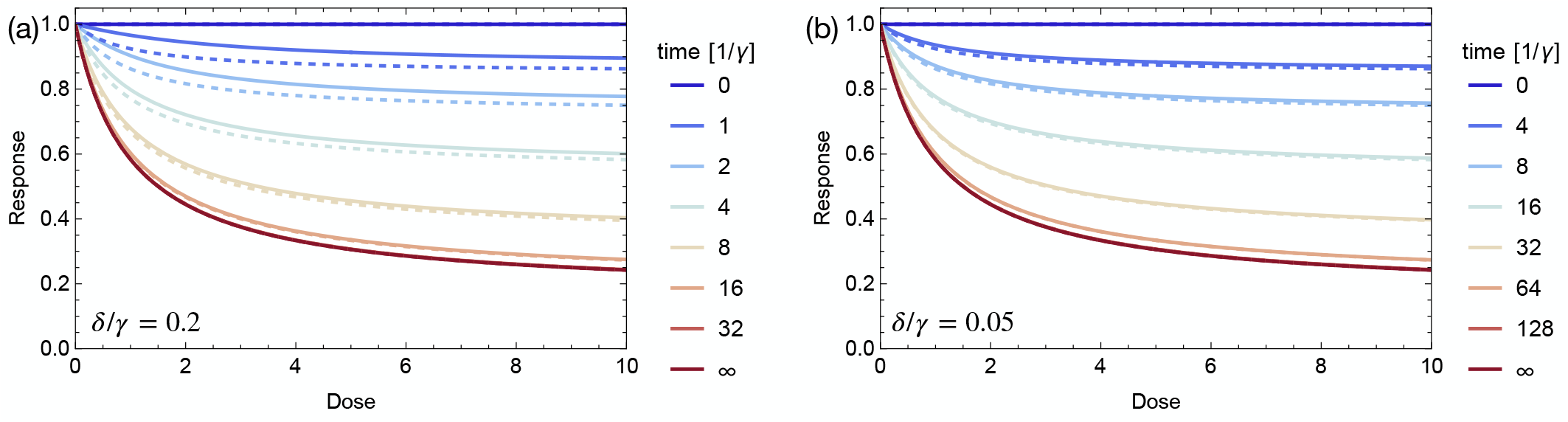
Response as a function of dose at various times after treatment in the regime that mRNA decay rate is much smaller than mRNA maturation rate: *δ ≪ γ*. In panel (a) *δ/γ* = 0.2 and in panel (b) *δ/γ* = 0.05. Solid lines correspond to exact solutions Eq. (5) while dashed lines correspond to approximate solutions Eq. (9). Observe that the response saturates with dose and approaches the steady-state solution at long times. Also observe that the discrepancy between exact and approximate solution decreases as the ration *δ/γ* becomes smaller. [to produce this plot we used *α/γ* = 1, *β/γ* = 1, and *κ/γ* = 5]

## III. THE DORESEQ NOISE MODEL

In order to connect the dose response model to the noise in gene expression, we introduce the following expression for the expected mean number of counts of gene *i* in sample *α*

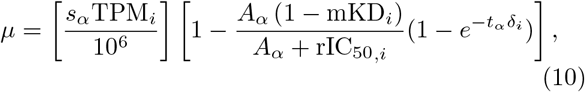

where *s*_*α*_ is the scaling parameter for sample *α*; *A*_*α*_ is the ASO dose in sample *α*; *t*_*α*_ is the time at which sample *α* was read out; TPM_*i*_ is the zero-dose transcripts per million value for gene *i*; mKD_*i*_, rIC_50,*i*_, and *δ*_*i*_ are the genespecific DoReSeq parameters as described in the previous section. In our analysis we set *s*_*α*_ = *N*_*α*_ to be the total number of non-duplicate reads of all genes in sample *α*, although other workers have used alternative normalization schemes [1, 2]. The term in the first bracket on the RHS of Eq. (10) is the expected number of counts for gene *i* at zero dose, while the term in the second bracket is the fractional decrease in the counts induced by the treatment. In the following, we will treat *s*_*α*_, *A*_*α*_, and *t*_*α*_ as confounding/independent variables, and TPM_*i*_, mKD_*i*_, rIC_50,*i*_, and *δ*_*i*_ as fit parameters.

We use the Negative Binomial distribution to describe the noise in NGS data, where the noise can be both biological or technical in nature. To simplify notation, we will parameterize the Negative Binomial distribution *P*_NB_(*x*|*µ, r*) using the mean, *µ*, and the dispersion parameter, *ϕ*. This parametrization is connected to the more standard parametrization in terms of *p* and *r* via

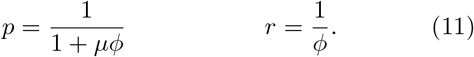

With our parametrization the mean and variance of the negative binomial distribution are

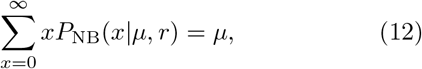

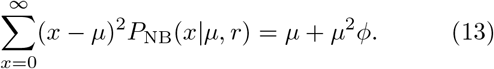

Next, we introduce a model for noise in the number of reads. This is the central equation of this paper:

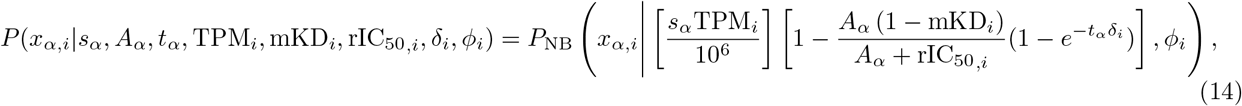

where *x*_*α,i*_ is the number of reads of gene *i* in sample *α* and *ϕ*_*i*_ is the gene-specific dispersion parameter. In other words, Eq. (14) says that that the distribution of *x*_*α,i*_ is the negative binomial distribution with mean set by the expected number of reads from Eq. (10) and width set by the gene specific dispersion parameter *ϕ*_*i*_. Further, in order to determine the expected number of reads, we need to have the values of the confounding/independent variables *s*_*α*_, *A*_*α*_, and *t*_*α*_ for sample *α* which are derived from the experimental data as well as the values of the fitting parameters TPM_*i*_, mKD_*i*_, rIC_50,*i*_, and *δ*_*i*_. We note that for the case where all measurements were carried out at the same time after treatment, the fitting parameter *δ*_*i*_ can be dropped and the expression 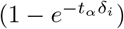 set to unity.

### IV. DORESEQ FITTING USING BAYESIAN INFERENCE

In this section we will specify our scheme for fitting dose response using Bayesian inference. Our goal is given dose-dependent experimental data to find the probability distribution of the fit parameters ℱ_*i*_ = {TPM_*i*_, mKD_*i*_, rIC_50,*i*_, *δ*_*i*_, *ϕ*_*i*_}

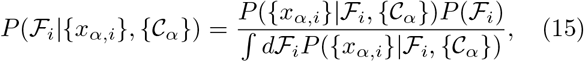

where {*C*_*α*_} = {*s*_*α*_, dose_*α*_, *t*_*α*_} is the set of confounding/independent variables. We use Eq. (14) to define the likelihood

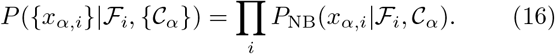

For the prior *P* (ℱ), we use box distributions with reasonable range for each of the fit parameters

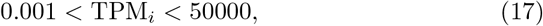

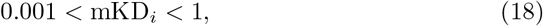

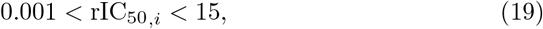

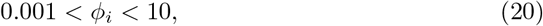

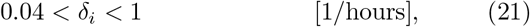

where *δ*_*i*_ is the only quantity with units and we measure it in inverse hours as indicated.

In order to efficiently explore the five dimensional space of fit parameters (four dimensional in case we do not have time-dependent data) we use Markov chain Monte Carlo (MCMC) to sample the numerator of the RHS of Eq. (15). The output of MCMC is a list of samples: 5-dimensional (4-dimensional if there is no time-dependence) vectors that define a set of models. The vectors sample the probability density defined by Eq. (15) – there are more vectors clustered in the regions where *P* (ℱ_*i*_|{*x*_*α,i*_}, {*C*_*α*_}) is large and fewer in the regions where it is small. We perform the sampling using the emcee python package [20].

To turn the list of vectors sampled by MCMC into a probability density we need to generate a normalized histogram. We focus on constructing three types of histograms: (1) histograms of the parameters that specify the noise distribution ℱ_DS_ = {TPM_*i*_, *ϕ*_*i*_}; (2) histograms of the dose response parameters ℱ_DR_ = {mKD_*i*_, rIC_50,*i*_, *δ*_*i*_} for the case we have time-dependent data or ℱ_DR_ = {mKD_*i*_, rIC_50,*i*_} for the case we only have data at a single time-point; (3) histograms of the observable KD_dose,time_, the knockdown at fixed dose and time. We use the histograms to obtain the maximum *a posteriori* (MAP) estimates of the fit parameters, credible regions [29] containing 50%, 95%, and 99% of the probability, and p-values for KD_dose,time_ to be above a threshold. We now comment on the three types of analysis that we perform.

The noise distribution histograms provide us with an estimate of the fit parameters ℱ_DS_ = {TPM_*i*_, *ϕ*_*i*_}, which are crucial for identifying poorly-behaved genes, i.e. genes that have a low number of reads (e.g. TPM_*i*_ *<* 10) or large variability (e.g. *ϕ*_*i*_ *>* 0.2). This is especially important with NGS data which includes thousands of genes (roughly 25,000 for our experimental DGE dataset) where it is infeasible to manually process the the data gene-by-gene. Additionally, noise distribution histograms are helpful for interpreting dose-response analysis — they provides a basis for understanding why certain genes have larger credible intervals than others when fitting dose response.

The dose-response histograms provide us with the data for quantifying the crucial property that we want to know: the dose-response parameters. Below, we show that the probability density function *P* (ℱ) is not separable, and therefore it is important to analyze it as a joint probability distribution. Further, we show that *P* (ℱ) contains a *sloppy* direction, which makes it hard to constrain the value of mKD_*i*_ and rIC_50,*i*_ simultaneously for weakly dose-responsive genes.

Finally, the fixed-dose, fixed-time response histograms combine the three dose-response fit parameters ℱ_DR_ = {mKD_*i*_, rIC_50,*i*_, *δ*_*i*_} into a single, biologically significant observable KD_dose,time_. KD_dose,time_ is helpful for predicting the outcome of a particular treatment: e.g. how much knockdown can one expect at at a fixed treatment dose? Further, we show that KD_dose,time_ is a *stiff* observable that makes it possible to identify strongly-, weakly-, and none-dose-responsive genes, e.g. using a p-value.

### A. Bayesian inference on synthetic DGE data

In this subsection we generate synthetic DGE data using the noise model of Eq. (14) and then use DoReSeq Bayesian inference to analyze it. The parameters that we use to generate synthetic data are typical of the fitted values for these parameters on experimental DGE data that we describe in the next subsection. The present subsection is structured as follows. First, we describe how the Bayesian inference machinery works by analyzing a small synthetic dataset with 3 genes, 102 samples at various doses, and no time dependence. Next, we generate 72 random replicates of our dataset and investigate replicate-to-replicate fluctuations. Then, we investigate how the sensitivity of our analysis depends on the number of samples, dispersion parameter, raw number of reads, and the total number of samples. We conclude this subsection by analyzing a synthetic dataset with time dependence.

#### 1. Setting up Bayesian inference machinery to fit synthetic data

We begin by considering a synthetic dataset that contains 102 samples and three genes. The dose values *A*_*α*_ for the samples consisted of

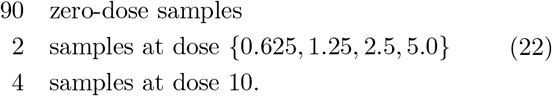

The scale factors *s*_*α*_ for the samples were chosen at random from a uniform distribution between 10^6^ and 10^7^. The three genes were chosen to contain a strongly dose-sensitive gene mKD_1_ = 0, a weakly dose-sensitive gene mKD_2_ = 0.5, and dose-insensitive gene mKD_3_ = 1.0. The ground-truth values for the other model parameters of the three genes were chosen to be identical: TPM_*i*_ = 50, *ϕ*_*i*_ = 0.05, and rIC_50,*i*_ = 2. To synthesize the number of raw reads *x*_*α,i*_ we dropped the time dependence from Eq. 14 to obtain

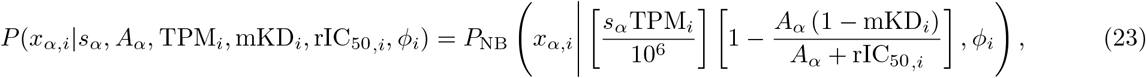

and then randomly sampled the negative binomial distributions. In order to visualize the simulated data, we converted the synthetic values of raw number of reads *x*_*α,i*_ to synthetic TPM (transcripts per million) using 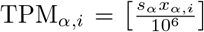, and plotted the results in Fig. 4(ac) using blue circles. We also plotted the ground truth TPM as a function of dose curves

**FIG. 4.**
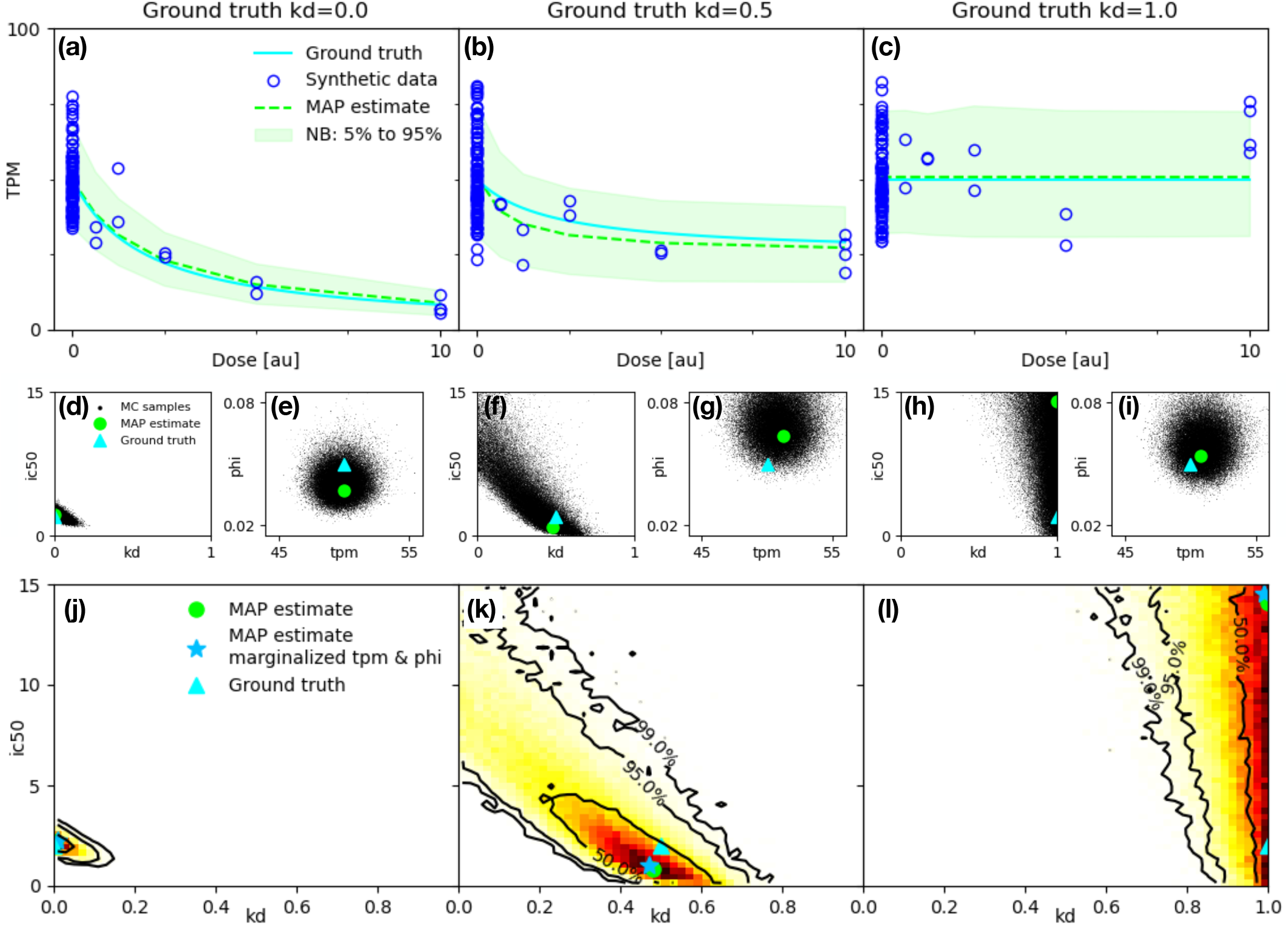
Fitting synthetic data with DoReSeq for a strongly dose responsive gene (left column), a weakly dose-responsive gene (middle column) and a dose non-responsive gene (right column). Panels (a-c): transcripts per million (TPM) vs. dose plots showing the ground truth curves, the synthetic data, and the Maximum A Posteriori (MAP) estimates of TPM vs dose curves (dashed lines) and negative binomial dispersion (green shaded regions, see supplement for details). Panels (d-i): Markov Chain Monte Carlo samples of the fitting parameters along with global map estimates and ground truth values of the fitting parameters. Panels (j-l): marginalized joint probability distributions for the dose-response fitting parameters mKD_*i*_ and rIC_50,*i*_. The global and marginalized MAP estimates of mKD_*i*_ and rIC_50,*i*_ are indicated along with the ground truth values. Black contours outline the credible intervals. (see text for details)

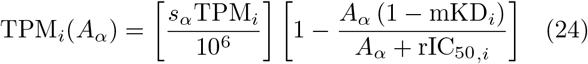

which appear in Eq. (23), for the three genes using cyan lines.

Next, for each gene *i* we sampled the distribution of the model fit parameters {TPM_*i*_, *ϕ*_*i*_, mKD_*i*_, rIC_50,*i*_}, given by Eq. (15), conditioned on the synthetic raw number of reads *x*_*α,i*_. Specifically, we fed *s*_*α*_, *A*_*α*_, and *x*_*α,i*_, along with the (un-normalized) probability defined by the numerator of Eq. (15), to the emcee Markov Chain Monte Carlo (MCMC) code. The code was setup to use 40 walkers. For each walker the first 1000 points were burned and the next 4000 points were recorded in order to obtain a total of 160,000 fit parameter samples for each gene. Here, each sample is a 4-vector composed of {TPM_*i,µ*_, *ϕ*_*i,µ*_, mKD_*i,µ*_, rIC_50,*i,µ*_}, where *µ* = 1, 2…160, 000 is the MCMC sample index.

We visualize the MCMC samples in Fig. 4(d)-(i). We begin by focusing on Fig. 4(d) and (e), in which we plot the MCMC sample for the gene with the strongest dose response. We also plot the ground truth value (i.e. the value of parameters we used to synthesize the data) as well as the global MAP estimate (i.e. the MCMC sample with the highest probability, see Eq. (15)). Fig. 4(f) and (g) pertain to the weakly dose-responsive gene while Fig. 4(h) and (i) pertain to the dose non-responsive gene. We now make several observations about the MCMC sampled distributions of fit parameters.

First, we observe that for all three genes TPM_*i*_ and *ϕ*_*i*_ are essentially independent of each other (see Fig. 4(e,g,i)), while mKD_*i*_ and rIC_50,*i*_ are strongly anti-correlated for the two dose responsive genes (see Fig. 4(d,f)). For the dose non-responsive gene, because mKD_*i*_ ≈ 1, the value of rIC_50,*i*_ does not strongly affect the shape of the TPM vs dose curve and therefore we observe that the distribution of rIC_50,*i*_ almost matches the prior, being uniformly distributed between 0.001 and 15 (see Fig. 4(h). The anti-correlation between mKD_*i*_ and rIC_50,*i*_ implies the existence of a *sloppy* direction [21]. We identify this *sloppy* direction as a simultaneous stretching out and lowering of the TPM vs dose curve by simultaneously increasing rIC_50,*i*_ and decreasing mKD_*i*_. Under this transformation the quality of the fit is largely unaffected because the TPM vs dose curve does not substantially change for small and intermediate doses. Comparing Fig. 4(d) and (f) we observe that the *sloppy* direction is significantly bigger problem for weakly dose-responsive genes. We stop the discussion of the *sloppy* direction for now, and come back to it later in the manuscript.

Second, we observe that the fractional variance of MCMC samples of TPM_*i*_’s is much smaller than that of *ϕ*_*i*_’s (see Fig. 4(e,g,i)). The underlying reason for this discrepancy is that it is much easier to estimate the mean of the negative binomial distribution as compared to its dispersion [30]. Indeed, DESeq2 package recommends using a bootstrapping strategy to address this issue, in which the distribution of *ϕ*_*i*_ values from genes of similar TPM’s is used as a prior when fitting *ϕ*_*i*_’s [2]. We are primarily interested in estimating knockdown as opposed to putting a tight bound on the dispersion parameter *ϕ*_*i*_ and therefore we simply fit *ϕ*_*i*_ on a gene-by-gene basis. However, in our case this fitting is strongly enhanced by having a large number (110) of zero-dose samples in our DGE datasets.

Let us focus on the gene knockdown parameters mKD_*i*_ and rIC_50,*i*_ next. These are the key parameters of interest as they encode the dose response of the gene. We, therefore, treat TPM_*i*_ and *ϕ*_*i*_ as nuisance parameters and marginalize over them. In Figs. 4(j-l), we plot the joint probability distributions for mKD_*i*_ and rIC_50,*i*_ which we obtain by histogram of the MCMC data. From the joint probability distributions we read off the marginalized MAP estimates for mKD_*i*_ and rIC_50,*i*_ (marked with a star in Figs. 4(j-l)) as well as the credible regions (the boundaries of the 50%, 95%, and 99% credible regions are marked with black lines in Figs. 4(j-l)).

### 2. Gene knockdown at a fixed dose: using Bayesian machinery on an observable

A useful alternative to estimating the fitting parameters is estimating the value of an observable. The observable that we focus on is the knockdown at a fixed dose. This observable is valuable for directly comparing the knockdown of different genes (e.g. to compare knockdown of off-target genes relative to the on-target genes). In Fig. 5, we show how we sample the probability distribution of knockdown at dose=5. We take the set of MCMC samples of {mKD_*i*_, rIC_50,*i*_}_*µ*_ (where *µ* is the MCMC sample index) and for each sample *µ* we compute the knockdown at dose = *A* = 5:

**FIG. 5.**
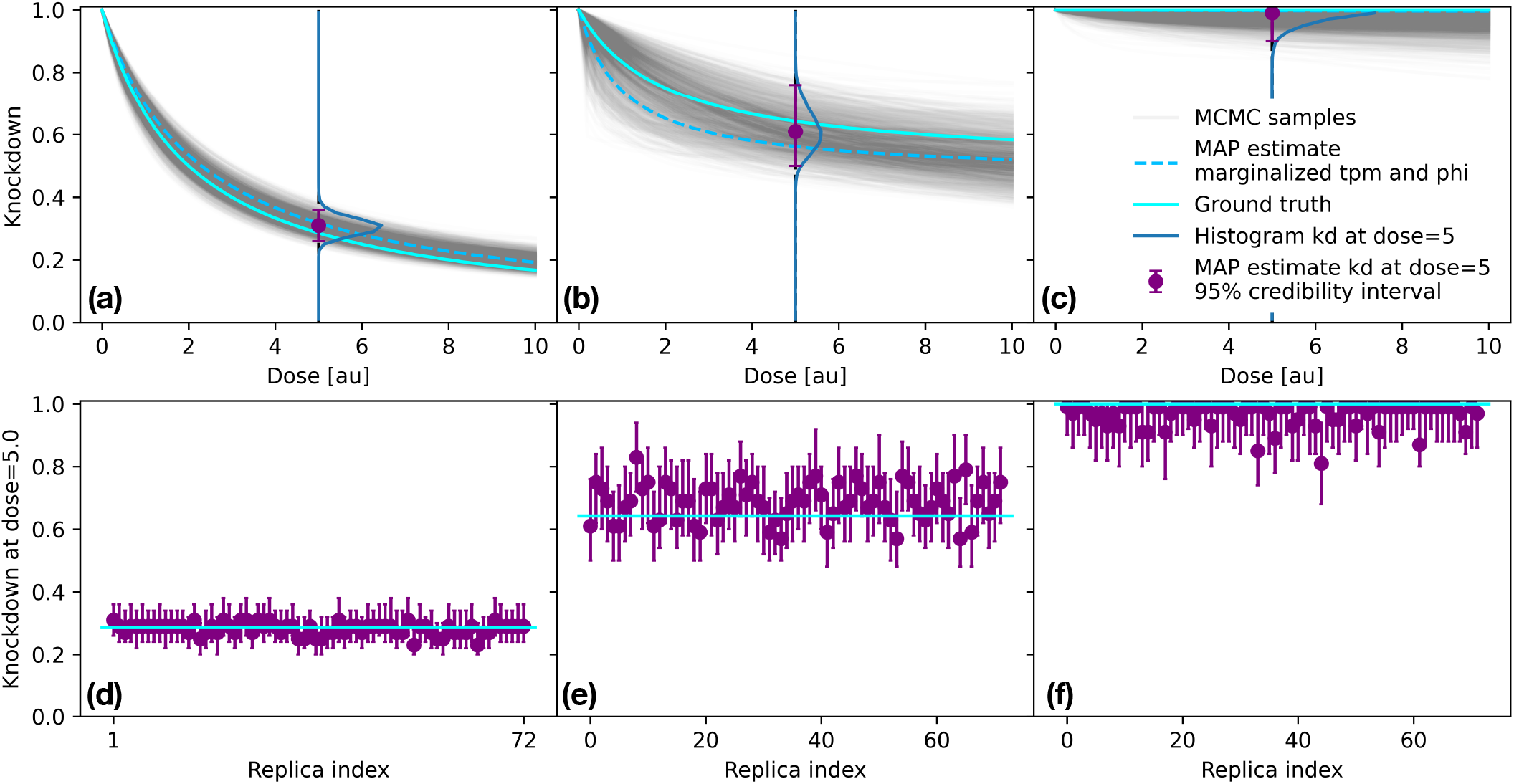
Estimating knockdown at a fixed dose. Panels (a-c): Knockdown as a function of dose. The gray lines represent a subsample of Monte Carlo data (showing 10,000 out of 160,000 samples). The MAP estimates of mKD_*i*_ and rIC_50,*i*_ marginalized over TPM_*i*_ and *ϕ*_*i*_ are shown with dashed light blue lines, while the ground truth curves are shown with solid cyan lines. The distribution of knockdown at dose 5 is shown with the solid blue lines, the MAP estimates and 95% credible intervals derived from these distribution are shown with the purple dots and errorbars. Panel (d-f): MAP estimates and 95% credible intervals over 72 replicates of the numerical experiment.

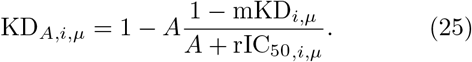

Next, we histogram KD_*A,i,µ*_ to obtain *P*_KD,*A*=5,*i*_(KD_5,*i*_) the distribution of knockdown at dose=5. From this distribution we find the MAP estimate of KD_*A,i*_ and the 95% credible interval (see Fig. 5(a-c)). Finally we repeat the procedure on 72 replicas of the numerical experiment and plot the resulting MAP estimates of KD_5,*i*_ and the 95% credible intervals in Fig. 5(d-f). We observe that that the credible intervals are only a little wider for the weakly dose responsive gene (Fig. 5(e)) as compared to the strongly dose-responsive and the dose non-responsive genes (Fig. 5(d) & (f)). At the same time, as expected, in almost all cases the 95% credible intervals straddle the ground truth.

We observe that the knockdown at dose=5 credible intervals (Fig. 5(d-f)) are much narrower than the credible regions of the maximum knockdown (Fig. 4(j-l)). Similarly, the spread of MAP estimates of knockdown at dose=5 is much narrower than that of the maximum knockdown (see Fig. 6 and supplement for additional details). This observation implies that KD_5,*i*_ lies largely along the stiff direction of the {mKD_*i*_, rIC_50,*i*_} space and therefore makes it feasible and allows us to recommend using response at a fixed dose as a way to identify dose-responsive genes. To help identify dose-responsive genes, we make one further step: we introduce a p-value for response of gene *i* at dose *A*. Specifically, the p-value measures the probability that KD_*A,i*_ is above a threshold *θ*

**FIG. 6.**
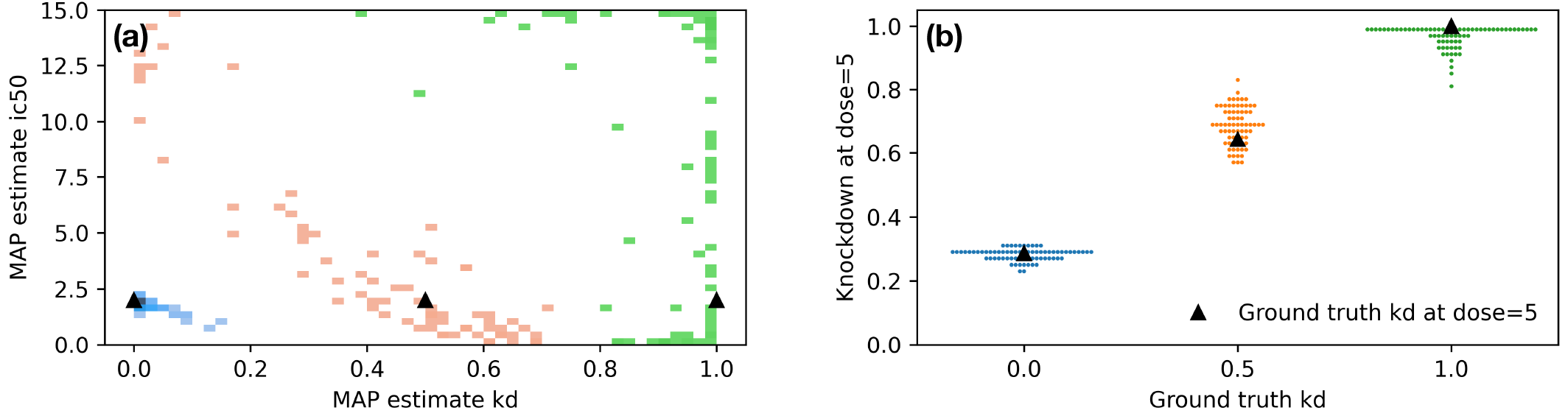
Comparing the dispersion of MAP estimates of mKD and rIC_50_ (Panel a) to MAP estimates of KD5 (Panel b) for synthetic data generated using three values of ground truth mKD: 0 (blue), 0.5 (orange), and 1 (green).

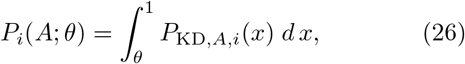

where a reasonable value for *θ* is 0.8. If the p-value for a certain gene is large, e.g. *P*_*i*_(*A* = 5; *θ* = 0.8) *>* 0.99, then it is very likely that the gene is not being knocked down. On the other hand, if the p-value is small, e.g. *P*_*i*_(*A* = 5; *θ* = 0.8) *<* 0.01, then it is very likely that the gene is being knocked down.

#### 3. Dependence of fit quality on parameters used to generate synthetic data

Here, we address how the fit quality depends on the parameters used to generate the synthetic data. In order to quantify the fit quality, we generate 288 replicates at each condition and compute *P*_KD,*A*=5,*i*_, the distribution of MAP estimates of knockdown at dose=5, as well as 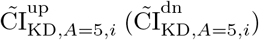, the median of the upper (lower) edge of the 95% credible intervals. We perform four separate sweeps in which we vary the ground truth value of one of the fit parameter at a time: mKD_*i*_, rIC_50,*i*_, *ϕ*_*i*_ and TPM_*i*_ while the remaining parameters are held fixed, see Fig. 5(a-d). We also perform a sweep of the number of samples while holding the ground truth values of all the other DoReSeq parameters fixed, see Fig. 5(e). When sweeping the ground truth values of the fit parameters, each replicate consisting of 102 samples with doses prescribed by Eq. (22) and scale factors *s*_*α*_ chosen from a uniform distribution between 10^6^ and 10^7^. On the other hand, when sweeping the number of samples, each replicate consists of an equal number of samples at each of the following doses: {0, 0.625, 1.25, 2.5, 5, 10} (e.g. when the number of samples is 12, there are 2 samples at each dose).

We observe several trends. First, as the mean expected number of raw reads becomes small at strong knockdown, the variance in the number of raw reads shrinks as prescribed by Eq. (13) and therefore we expect that the quality of fit will improve. We can observe the relationship between gene expression and the width of the negative binomial distribution in Fig. 4(a), where the green shaded region denoting the width of the negative binomial distribution becomes narrower as gene expression is knocked down. The relationship between stronger knockdown and higher quality fitting can be seen in Fig. 5(a) at low ground truth values of mKD_*i*_ as well as Fig. 5(b) at low ground truth values of rIC_50,*i*_.

Second, if there is very little knockdown, the quality of the fit will also improve because the prior encoded by Eq. (18) will squeeze the distribution of fitted knockdowns from above. This effect can is observed in Fig. 5(a) as the ground truth values of mKD_*i*_ → 1.

Third, for larger values of the ground truth dispersion parameter *ϕ*_*i*_ the synthetic data has larger variance and hence the quality of the fit degrades. This trend is observed in Fig. 5(c), where fitting becomes unreliable for ground truth *ϕ*_*i*_ ≳ 0.256.

Fourth, DGE is expected to lose precision when the raw read counts become small. This loss of precision is a property of the Negative Binomial distribution, in which the ratio of the standard deviation to the mean

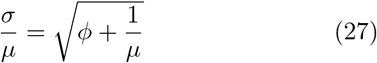

starts to grow rapidly for *µ* ≲ 1*/ϕ*. In our case, this condition starts to set in for TPM_*i*_ ∼ 2 − 20 (where the range is set by the distribution of scale factors 10^6^ ≤ *s*_*α*_ ≤ 10^7^). We indeed observe this trend in Fig. 5(d). We see a slow degradation of the quality of fit as the ground truth value decreases to TPM_*i*_ = 16. At TPM_*i*_ = 8 the fit becomes significantly worse, and for TPM_*i*_ ≤ 4 the fit becomes unreliable as the median value of the upper edge of the credible intervals sharply jumps up and the distribution of MAP estimates acquires weight at KD_5,*i*_ = 1.

Fifth, we expect that the fit quality degrades as the number of samples decreases. We indeed observe this trend in Fig. 5(e).

#### 4. Time dependence

In this subsection we add the time dimension to synthetic data. Specifically, we synthesize a dataset that contains samples measured at 2, 4, 8, and 16 hour marks. At the 2, 4, 8, and 16 hour marks the dataset contains two samples at dose {0.625, 1.25, 2.5, 5} and four samples at dose 10. In addition the dataset contains 90 zero-dose samples. As before, we chose the scale factors for each sample at random from a uniform distribution between 10^6^ and 10^7^. We generated synthetic data for four genes with *δ*_*i*_ = {0.5, 0.25, 0.12, 0.6} [1*/*hours] and the ground truth values for the remaining fit parameters set to TPM_*i*_ = 50, *ϕ*_*i*_ = 0.05, mKD_*i*_ = 0, and rIC_50,*i*_ = 2. In Fig. 8(a-d) we plot the time-dependent synthetic data that we are going to fit with points and the corresponding ground-truth TPM vs. dose curves with solid lines.

Next, we applied our Bayesian inference machinery to fit the time dependent synthetic data. To asses the quality of the fit, we plot the TPM vs. Dose curves obtained from the global MAP estimate of the fit parameters in Fig. 8(a-d) with dashed lines. We observe that in all cases the fits are quite close to the ground truth curves (solid lines). Further, in Fig. 8(e-h), we plot the MCMC generated distributions of the fit parameter *δ* (i.e. the RNA decay rate) for the four different ground truth values of *δ*. From the MCMC distributions we also obtain the marginalized MAP estimates for *δ* as well as the 95% credible intervals. We observe that the MAP estimates (listed in the figure) are close to the ground truth values of *δ* and the credible intervals straddle the ground truth. In summary, the evidence we presented in this subsection indicates that it should be possible to extract the RNA decay rate from experimental data. It is important to have sufficient quantity and quality of DGE data and to choose the measurement times so that there is considerable variation in the expression of the target gene (that is the measurement times are dictated by ∼ 1*/δ*).

### B. Bayesian inference on experimental DGE data

The goal of this subsection is to apply the machinery that we have developed to analyze an experimental DGE dataset. We aim to both identify dose-responsive genes and quantify their response.

The dataset we analyze consists of 121 samples, with 110 zero-dose samples, 2 samples at dose {0.625, 1.25, 5}, 1 sample at dose {2.5}, and 4 samples at dose {10}— all ASO doses are in micro-molar (*µ*M) units. The total number of non-duplicate counts ranged from 0.49 to 8.93 million with an average of 3.22 million per sample. We selected 8005 gene, with zero-dose TPM_*i*_ ≳ 10, for analysis. As the experimental dataset we analyze is proprietary, we are unable to publish the ASO sequence nor the gene labels. We have included the table of raw counts and the table of metadata in the supplementary materials. The python code that we wrote to analyze the data and produce the figures i n t his m anuscript a nd i n the supplement is available for download.

In order to remove unwanted variance from plateto-plate variations, we note that when applying our Bayesian machinery to experimental data we used a separate and independent TPM parameter for each plate. Specifically, we replace Eq. (14) with

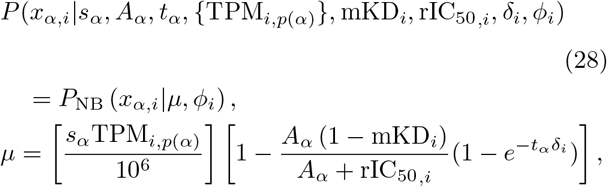

where {TPM_*i,p*(*α*)_} is a collection of TPM values associated with gene *i*, with one value assigned to each plate *p*(*α*). In our case there were 8 plates and hence 8 values of {TPM_*i,p*(*α*)_} were fitted for each gene. In order to compare experimentally obtained TPM values across different plates, we introduce a normalization scheme where we normalize the experimentally observed TPM value, on a plate-by-plate basis, to the mean of the MAP estimates of TPM over the plates

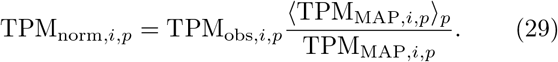

We also note that we implemented the mixed probability distribution method for attenuating outliers. In this method, we supplement the Negative Binomial distribution for inliers with a much wider Negative Binomial distribution for outliers. For our experimental dataset, we found that attenuating the outliers was not particularly important and hence the data analysis that we report here has this feature turned off. However it is available in our *DoReSeq* code should other researchers be interested in using it.

In order to identify dose-responsive genes, we computed the p-value of the knockdown at dose=5 for each of the 8005 genes. The p-value was evaluated using Eq. (26) with threshold set to *θ* = 0.8. The histogram of p-values, see Fig. 9, appears to be bimodal with a peak at small pvalues that corresponds to strongly dose-responsive genes and another peak at large p-values that corresponds to the dose-non-responsive genes.

Next, in order to better understand the characteristics of the genes in the two peaks of the p-value distribution, as well as genes in between, we choose three representative genes with TPM ∼ 50, one strongly dose-responsive gene (p-value=0), one weakly responsive gene (p-value=0.123) and one dose non-responsive gene (pvalue=1). We plot the results for all 203 strongly dose responsive genes and a samples of 100 weakly dose-responsive genes and 100 dose non-responsive genes in the supplement.

To visualize the dose response, we plot the normalized experimental TPM values (see Eq. 24) as a function of dose in Fig. 10(a-c). Superimposed on the experimental points are MAP estimates of TPM as a function of dose (Eq. 24 obtained using the plate-averaged value of TPM: TPM_*i*_ → ⟨TPM_MAP,*i,p*_⟩_*p*_) as well as the MAP estimate of the dispersion encoded as the 5% and 95% quantiles of the negative binomial distributions (see supplement for details). We observe that for the strongly dose-dependent gene, both the experimental data and the MAP estimate decrease substantially with dose (see Fig. 10(a)). On the other hand for the weakly dose-dependent gene, the decrease with dose is much less pronounced Fig. 10(b)) and for the dose-independent gene no decrease in gene expression with dose is observed Fig. 10(c)).

To understand the quality of the fit, we look at the probability distributions for the knockdown parameters (see Fig. 10(d-i)). We observe that for the strongly dose-responsive gene the MAP estimates of mKD and rIC_50_ come with reasonably tight credible regions (see Fig. 10(d)). For the weakly dose-responsive gene the credible regions are much less tight (see Fig. 10(f)). Finally for the dose-non-responsive gene the credible region is tight on mKD but loose on rIC_50_ (which is expected when there is no knockdown, see Fig. 10(i)). These observations are consistent with our analysis of synthetic data in subsection IV A 1. We observe tight credible intervals for knockdown at dose=5 for all three genes, see Fig. 10(e,g,i). This second observation is again consistent with our analysis of synthetic data, see discussion in Sec. IV A 2).

We argue that our observations on both synthetic and experimental data indicate that response at fixed dose is a reasonable way to identify dose-responsive genes (given good experimental design: appropriately chosen dosing, readout time, and number of samples). Genes that are indeed dose responsive can be further analyzed by breaking down the response into parameters mKD and rIC_50_.

## V. DISCUSSION

We begin by commenting on the connection between DoReSeq kinetics and the Monod or the Hill equations. The Hill equation describes the equilibrium of a reversible chemical reaction that is completely unrelated to the kinetics that we describe in DoReSeq. In contrast, the Monod equation describes the *rate* of bacterial growth in the presence of a substrate limiting growth. The fact that in the long-time, *steady-state* limit DoReSeq doseresponse curve has the functional form of a Monod equation is curious, though, the variable in the functional form for DoReSeq is a *concentration* as opposed to a *rate* in the Monod equation. Both DoReSeq and the Monod equation describe kinetics, although the details of the kinetics and mechanisms are completely different, e.g. in case of DoReSeq the rate of RNA production is maximal in the absence of ASO, while the rate of bacteria growth is zero in the absence of the rate limiting substrate in the Monod equation. We belabor these points to distinguish our work from the phenomenological methods of steadystate dose response fitting commonly used in ASO and siRNA fields. Also, our motivation in this work was not to just fit steady state dose-response but build a model for time-dependent dose-response kinetics that includes target-RNA half-life.

One of the key feature of DoReSeq kinetic model is capturing the “window-of-opportunity” phenomena—the maximum knockdown of gene expression that an ASO can achieve is not controlled by the concentration of the ASO alone, but rather by the competition between the maturation of the pre-mRNA and the rate at which the ASO-bound pre-mRNA is cleaved by RNase H1. Further, from our kinetic model we find that the concentration of the mature mRNA exponentially decays in time to its long-time value, with the rate constant set by the mRNA decay rate.

Being able to compute the credible intervals provides drug developers in oligonucleotide space with improved quantification for evidence-based decision making on what makes a specific ASO or siRNA—for example, a time-course in DGE analysis across multiple doses can be used to measure on-target response and identify offtargets in parallel. For off-targets, the time course informs us on primary hybridization-mediated off-targets and cytotoxicity-mediated off-target (pathway modulation). Time is an important confounding variable. DoReSeq includes a model of kinetics and thus allows us to fundamentally analyze the time dependence of gene expression in response to dosing. Experimentally, capturing time dependence purely from DGE dose-time course is somewhat challenging because we need sizeable dose-time samples to constrain the RNA decay rate parameter *δ*. However, decay rates can be learned transcriptome-wide in other assays [31]. Using synthetic data, we performed a numerical experiment to show that DoReSeq is indeed able to capture time dependence for genes that show strong dose-response. In this paper, we do not present experimental data across multiple time points, but will do so in future work. The close agreement of DoReSeq analysis of synthetic and experimental single-timepoint data gives us confidence in our methods.

In this work, we have focused on gene knockdown. However, the Baeysian statistical approach is general and can be used with different, perhaps phenomenological, dose-response models. For example, we can *ad hoc* extend the dose-response model to gene up-regulation by allowing for *mKD >* 1, deviating from the theme of this paper of deriving the underlying kinetics. Moreover, though we motivated the dose-response model with RnaseH-engaging ASO-mediated knockdown of RNA, it is far more general. Such generality is not uncommon in system-level modeling [32]. Fundamentally, the model captures a “window of opportunity”— the target loci in the RNA is available for enzymatic activity for knockdown effects only within a limited timescale, and kinetic race between the events of competing regulatory processing of RNA and enzymatic activity of knockdown mechanism-of-action are important. For ASOs, we have argued that co-transcriptional events like exon-definition and splicing, RNA-binding protein (RBP) occupancy, RNA-secondary etc. are reasonable mechanisms by which the target loci is rendered less “targetable”. For siRNAs, events after nuclear to cytoplasmic transport—for example, RBP and translational machinery engagement, regulatory-RNA interactions etc. can create a similar window-of-opportunity. The model does not need to commit to any one mechanism in particular, but if the timescales of any such events compete with the rate at which RNAi-machinery can knockdown mature RNAs, empirically, targetability constraints will be observed for siRNAs too [33]. We ignore the complexity of different timescales that may be involved for exon-targeting ASOs [23]. These ASOs can act both co-transcriptionally and in the cytoplasm on mature RNAs. Modeling this effect will need more parameters corresponding to the fate of the pre-mRNA and mature mRNA, and is left to future work.

Another advantage of our work is that, unlike DESeq/DESeq2 methods, we use the full power of Bayesian inference. Instead of just identifying MLE/MAP points, we compute the whole distribution. This allows us to construct p-values and credible intervals that directly quantify how constrained our fit is by the available data. The additional computational cost for our method is reasonable, e.g., about 2 hours to process 8000 genes on a 72-core machine.

Besides computational cost the experimental cost is substantially higher compared to standard practice because we need more samples across multiple doses— however we believe this is necessary for creating safer OBMs. The benefits of DoReSeq is not obvious if the experimental design involved just two doses; an untreated control sample-set in replicates and a single high-dose treated sample-set in replicates. To take advantage of DoReSeq, dose-response should ideally be sampled at multiple-doses with replicates at each dose.

Finally we point out that DoReSeq can be used to study experimental design. For example by fitting synthetic datasets with different number of samples, e.g. see Fig. 7(e), one can figure out the statistical power of the experimental design (dosing regimen, number of replicates, depth of sequencing etc.) to detect and quantify dose-response reliably. The optimization of Design of Experiments given a fixed budget of samples and sequencing is left for future work.

**FIG. 7.**
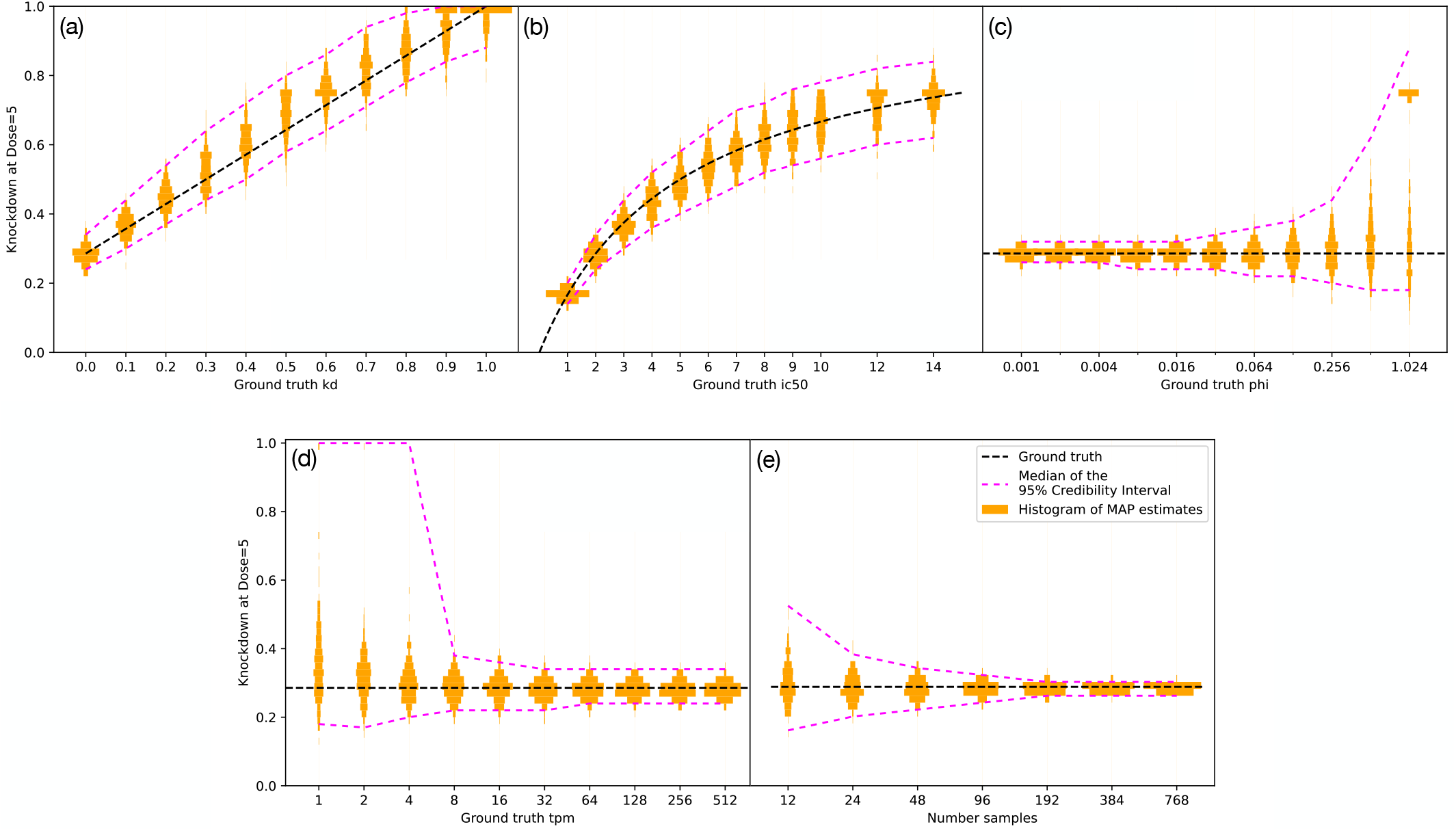
Effect of varying parameters on fitting synthetic data. Panels (a)-(e) show histograms of MAP estimates of Knockdown at Dose=5 obtained on synthetic data as well as the ground truth expectation of the knockdown. Panels (a)-(d): we fixed three of the four fit parameters and tuned the value of the remaining fit parameter. The values of the tuned parameter are indicated by tick marks on the horizontal axis (the remaining parameters were fixed to mKD = 0, rIC_50_ = 2, *ϕ* = 0.05, TPM = 50). For each value of the tuned parameter we generated and fitted 288 realizations. We observe that the spread of MAP estimates of knockdown at Dose=5 is weakly dependent on the ground truth values of knockdown mKD and rIC_50_. On the other hand, the spread of MAP estimates increases with the dispersion parameter *ϕ*. It also increases as the number of raw reads becomes small, which we control by setting the ground truth value of the TPM parameter to be small. Panel (e): we fixed mKD = 0, rIC_50_ = 2, *ϕ* = 0.05, TPM = 50 and varied the total number of synthesized samples that we fit. We observe that as the number of samples grows the spread of MAP estimates of knockdown at dose=5 decreases.

**FIG. 8.**
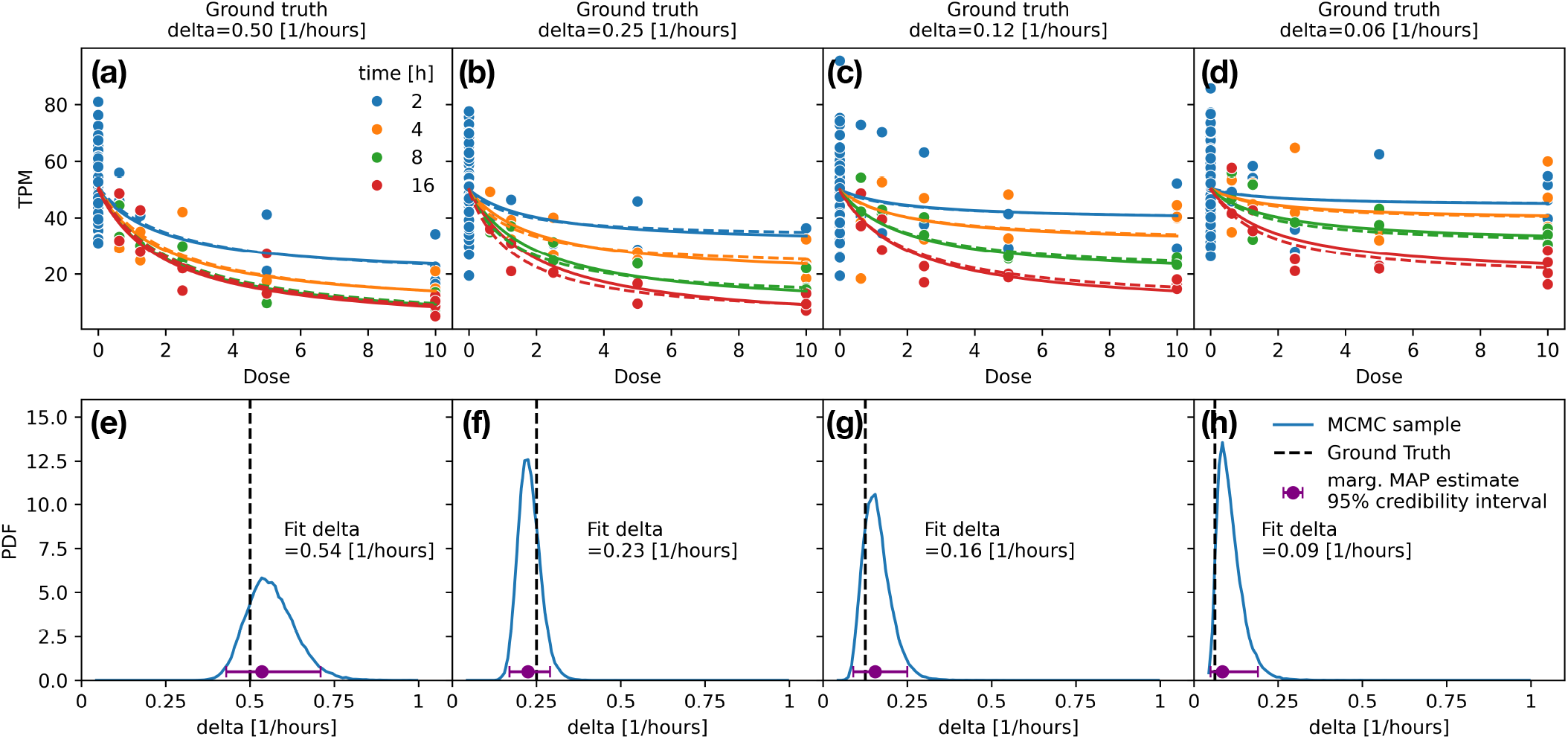
Fitting the RNA decay rate parameter *δ* with time-dependent data. Panel (a-d): TPM vs dose plots for four different values of *δ* (as indicated in the column titles). Points represent synthetic data for four different times (as indicated in the legend). Solid lines represent the ground truth TPM vs. dose curves and dashed lines represent the fitted TPM vs. dose curves (with parameters obtained from the global MAP estimate). Panel (e-h): marginalized probability distributions of *δ* obtained from the Markov Chain Monte Carlo. Ground truth value of *δ* is indicated by the dashed black line in each panel. The marginalized MAP estimate and the 95% credible interval for *δ*, obtained from the probability distribution, is indicated by the purple dot and bars in each panel.

**FIG. 9.**
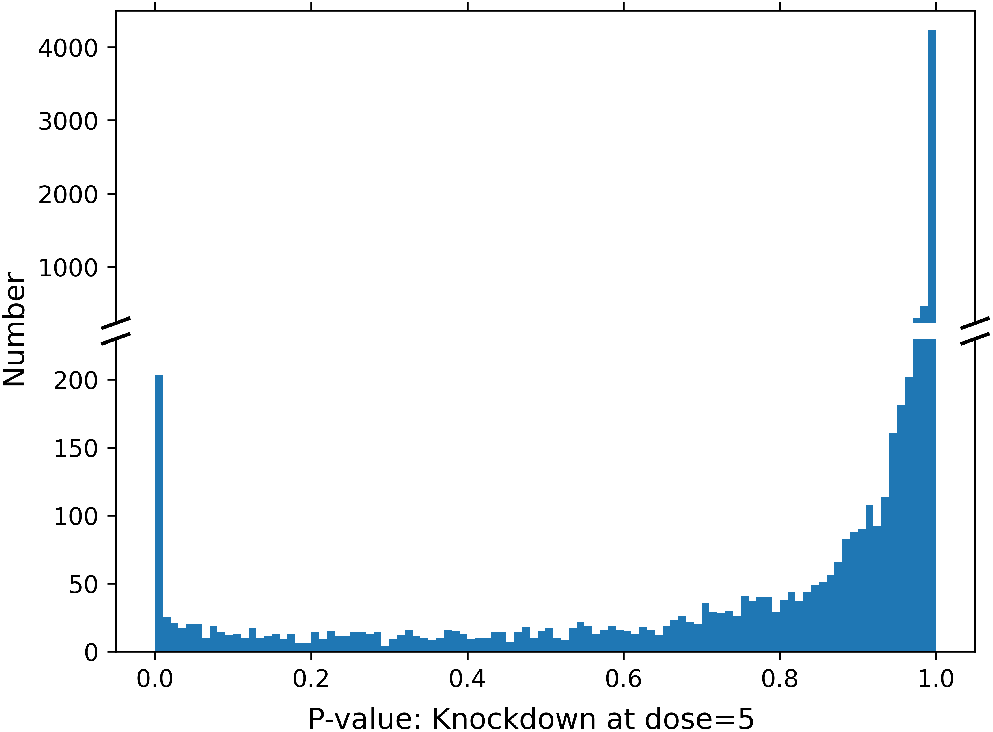
Histogram of knockdown at dose=5 p-values computed for experimental data. The histogram shows 203 genes that are strongly knocked down (p-value¡0.01), 3573 genes that are weakly knocked down (0.01¡P-value¡0.99), and 4229 genes that are not knocked-down (0.99¡ p-value). The vertical axis is split in order to show the peak at high P-values that corresponds to genes that are not knocked down.

**FIG. 10.**
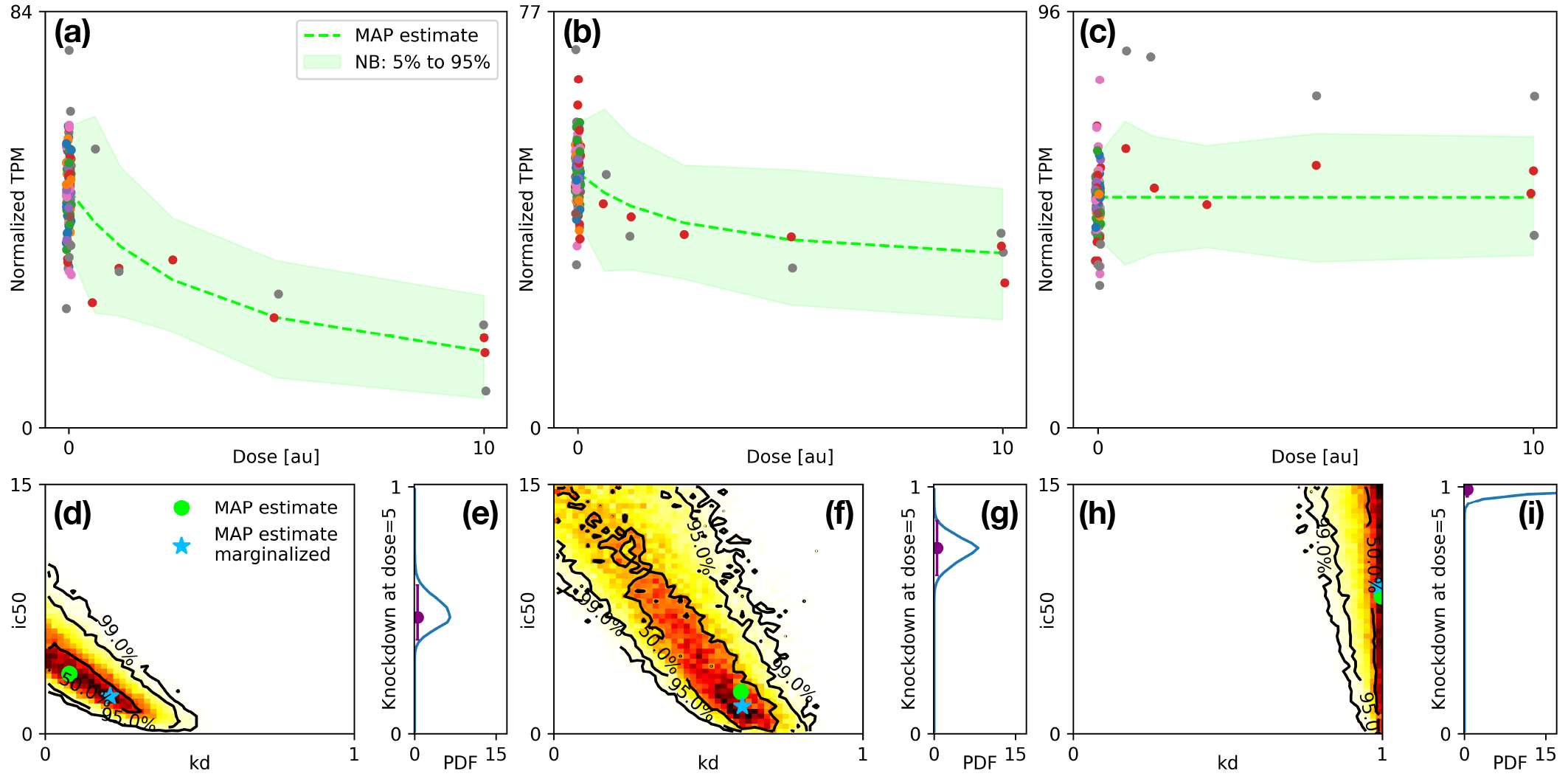
Bayesian inference fits of experimentally observed dose-response on a typical strongly dose-responsive gene (left column), weakly dose-responsive gene (middle column) and a dose-non-responsive gene (right column). Panels (a,b,c) show the experimentally observed dose response for the three genes using the colored points, where color corresponds to the eight plates that were processed to obtain this data. The experimental TPM values are regularized using MAP estimates of TPM on a plate-by-plate basis (see text). The global MAP estimate for dose dependence of TPM and width of negative binomial distributions are indicated with the green dashed line and the green shaded region. Panels (d,f,h) show the histogram for dose-response fit parameters mKD_*i*_ and rIC_50,*i*_. The global and marginalized MAP estimates are indicated by the dots and the 50, 95, and 99% credible regions by the black lines. Panels (e,g,i) show the histograms for the knockdown at dose=5 along with marginalized MAP estimates and 95% credible intervals indicated by the purple dots and bars.

## VI. CONCLUSION

In this paper, we described a method for fitting doseand time-dependence of gene expression as measured by DGE, or other similar Next Generation Sequencing (NGS) methods.

We presented a kinetic model for ASO and RNaseH mediated knockdown of gene expression. Using our kinetic model we derived a time-and dose-dependent parametrization for the response of gene expression to an ASO, Eq. 9. Next, we constructed a noise model for gene expression as measured by DGE (or other next generation sequencing methods). In our noise model, the number of reads of a particular gene is drawn from a Negative Binomial distribution with a timeand dosedependent mean and a gene-dependent dispersion parameter. Specifically, the mean is constructed by multiplying our timeand dose-dependent parametrization for RNA concentration by a scale factor (i.e. the total number of reads for all genes). In order to fit gene expression data, we use the machinery of Bayesian inference and Markov Chain Monte Carlo to construct the probability distribution of fit parameters conditioned by the experimental data.

We have tested our Bayesian inference machinery both on synthetic data as well as on experimental DGE data. In both cases we find that we can distinguish dose-responsive genes from dose-non-responsive genes using the P-value for response at fixed dose, given by Eq. (26). We identify genes that have low p-values as dose-responsive. For the dose-responsive genes, we can further characterize the dose response by finding the Maximum A Pasteriori (MAP) estimates of three parameters: the maximum knockdown mKD, the relative IC50 rIC_50_ and, if sufficient time-dependent data is available, the RNA decay rate *δ*. mKD and rIC_50_ are the crucial two parameters, beyond the certainty of dose-responsiveness, as they enable us to determine if the knockdown is clinically relevant at the dose that we will provide to set the on target effect. Along with MAP estimates we also compute credible regions for these parameters, precisely quantifying how well the experimental data measures the values of these parameters. We find that the credible regions for these parameters are much broader for weakly dose-responsive genes, and much sharper for strongly dose-responsive genes, owing to the presence of a *sloppy* direction in the {mKD, rIC_50_} space. We recommend using response at fixed dose in order to identify strongly-, weakly-, and dose-non-responsive genes (using p-values) because this observable is more constrained—it is essentially orthogonal to the *sloppy* direction associated with increasing rIC_50_ (the relative IC_50_) and decreasing mKD (the saturating knockdown value).

We hope our methods will be adopted for better quantification of onand off-target of OBMs. We believe that such an adoption will ultimately result in better engineered and safer OBMs, and avoid potentially dangerous off-target gene knockdown.

## ACKNOWLEDGMENTS

The authors thank Niu Du and Jaspreet Bhamra for useful discussions, and Christopher Hart for useful discussion, support and encouragement.

## Notes

### Competing Interest Statement

All authors work at Creyon Bio, a company that engineers Oligonucleotide Based Medicines.

