## Additional figures and SI for "A Dose-Response Model for Accurate Detection and Quantification of Transcriptome-Wide Gene Knockdown for Oligonucleotide-Based Medicines"

### Supplementary material: A Dose-Response Model for Accurate Detection and Quantification of Gene Knockdown with Next Generation Sequencing Data

David Pekker,* Steven Kuntz, Monica McArthur, Tim Nicholson-Shaw, Sara Yanke, and Swagatam Mukhopadhyay†
*Creyon Bio, Carlsbad, Ca 92010*

#### I. CORRELATIONS BETWEEN FIT PARAMETERS

In this section we investigate whether there are any correlations in the MCMC samples of fit parameters. Specifically, we re-plot the data of Fig. 4(d-i) of the main text using all possible pair-wise combinations of fit parameters (see Fig. S1). We observe that except for the correlation between mKD and $\mathrm{rIC}_{50}$ that was discussed in the main text, there are no strong pair-wise correaltions.

#### II. SLOPPY DIRECTIONS: MAP ESTIMATES OF mKD AND $\mathrm{rIC}_{50}$ VERSUS $\mathrm{KD}_5$.

In this section we compare how well we can estimate the dose response parameters maximum knockdown mKD and relative inhibitory concentration $\mathrm{rIC}_{50}$ versus how well we can estimate $\mathrm{KD}_5$ the knockdown at dose 5. Specifically, we generate synthetic data, with different total number of samples as well as different mKD, and investigate how well we can recover the various parameters.

We have simulating measuring three genes with $\mathrm{mKD}_i = \{0, 0.5, 1\}$, $\mathrm{rIC}_{50,i} = 2$, $\phi_i = 0.05$ and $\mathrm{tpm}_i = 50$ in 7 different experiments that utilize $12, 24, 48, ..., 768$ samples. For each experiment we considered six doses $\{0, 0.625, 1.25, 2.5, 5, 10\}$, with equal number of samples at each dose. Each of the 7 experiments has been replicated 288 times to gain statistics on how well we are estimating mKD, $\mathrm{rIC}_{50}$, and $\mathrm{KD}_5$.

We observe (see Fig. S2) that for the strongly dose-responsive gene (blue bars) as well as for the dose non-responsive gene (green bars) mKD and $\mathrm{rIC}_{50}$ estimates and $\mathrm{KD}_5$ estimates work well, even when the number of samples is very low. For the weakly dose-responsive gene (orange bars), $\mathrm{KD}_5$ estimates continue to work well even with few samples. However, mKD and $\mathrm{rIC}_{50}$ estimates for the weakly dose-responsive gene only work well when the number of samples is large, $\gtrsim 192$. Specifically, we see that when the number of samples is $\lesssim 192$, it is not possible to simultaneously determine both mKD and $\mathrm{rIC}_{50}$ due to a sloppy direction.

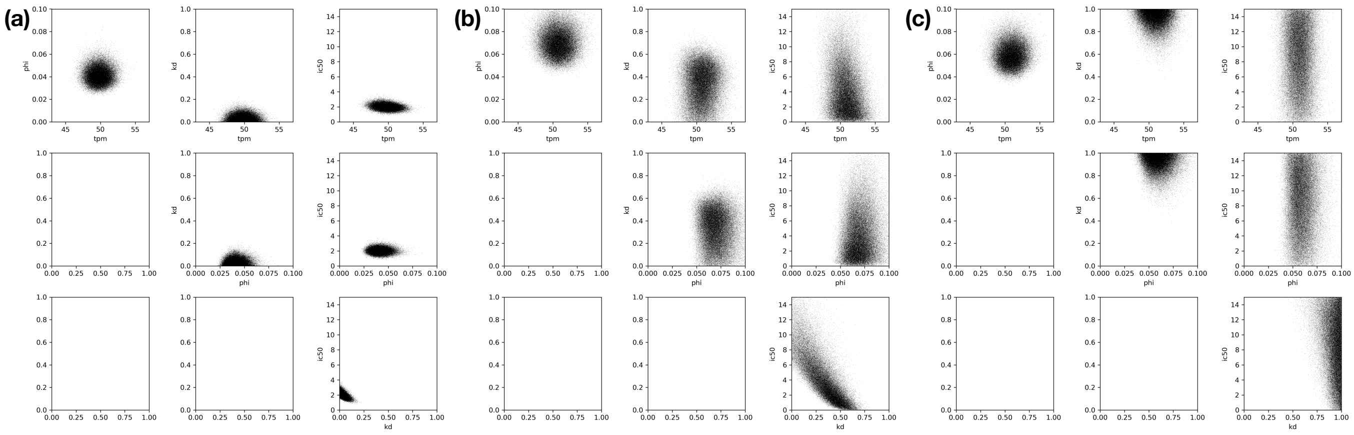


FIG. S1. Pairwise correlations between fit parameters in MCMC samples. Data is identical to the one displayed in Fig. 4(d-i) of the main text. Panel (a) corresponds to data from panels (d,e) of the main text, panel (b) to panels (f,g), panel (c) to panels (h,i).

*
†

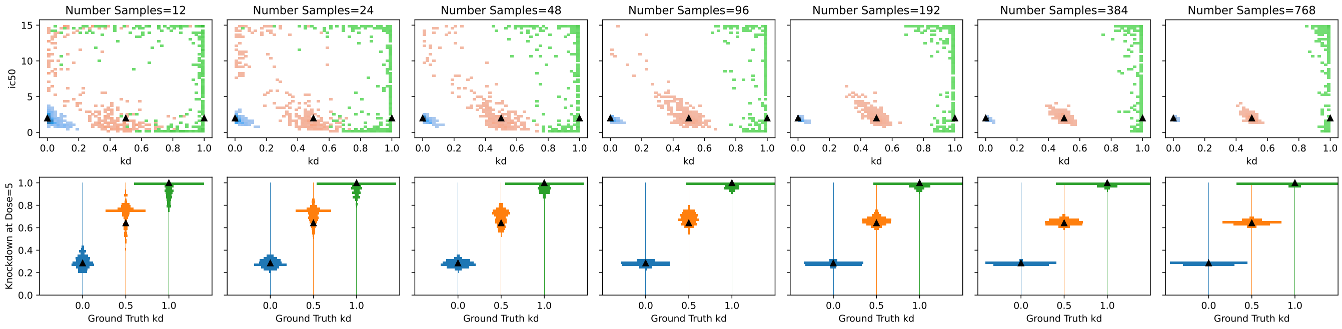


FIG. S2. Top row: histograms of marginalized MAP estimates of knockdown parameters $\mathrm{mKD}_i$ and $\mathrm{rIC}_{50,i}$, as a function of the number of samples. Bottom row: histograms of $\mathrm{KD}_5$, the knockdown at Dose=5, as a function of the number of samples. The ground truth values of mKD used to generate the data were 0.0 (blue bars), 0.5 (orange bars), 1.0 (green bars).

#### III. VISUALIZING THE MAP ESTIMATE OF THE NEGATIVE BINOMIAL DISPERSION

In this section we illustrate how we compute (and plot) the 5% and 95% quantiles of the negative binomial distributions in Fig. 4(a,b,c) and Fig. 10(a,b,c) of the main text.

We start from the global MAP estimate for gene $i$: the plate-dependent $\mathrm{tpm}_{i,p}$, the negative binomial dispersion parameter $\phi_i$, the maximum knockdown mKD, and the relative $\mathrm{rIC}_{50}$. Next, for each experimental sample $\alpha$, we construct 1000 random samples of raw read counts using Eq. 28 (or Eq. 14 if we do are not using plate-dependent tpm) of the main paper, using the experimental values of the independent/confounding parameters the scale factor $s_\alpha$, the dose $A_\alpha$, and the time $t_\alpha$ (if time-dependence is being used). This numerically generated data represents the dispersion of the MAP estimate of the fit parameters.

In Fig. S3(a-c), we plot the histograms of the numerically generated raw read counts, aggregated by plate and dose. While Fig. S3(a-c) is helpful for visualizing the numerically generated data, it is not helpful for interpreting it because different samples have wildly different scale factors $s_\alpha$.

In order to make it easier to compare different samples, we take our numerically generated raw read counts and convert them into transcripts per million (tpm). The histograms of the numerically generated tpm values, again aggregated by plate and dose, are plotted in Fig. S3(d-f). From Fig. S3(d-f), we observe that experimental data largely falls inside the numerically generated histograms, indicating that the MAP estimate of the dispersion parameter $\phi$ is reasonable.

In order to make it easier to compare samples on different plates, we normalized the tpm values to the mean over the plates (see Eq.  of 29 the main text). The resulting histograms are plotted in Fig. S3(g-i). Using normalized tpm values indeed helps to remove a lot of unwanted variance and makes it easier to compare samples on different plates.

Finally, we comment on the green shaded regions in Fig. 4(a,b,c) and Fig. 10(a,b,c) of the main text.

In Fig. 4(a,b,c) we were investigating synthetic data generated without plate-dependent tpm. Therefore, we numerically generated tpm values as described in this supplement and aggregated those values by dose. For each dose we computed the 5% and 95% quantiles of the numerically generated tpm values and plotted the results in Fig. 4(a,b,c) of the main text.

In Fig. 10(a,b,c) we were investigating the experimental data with plate-dependent tpm fit. Therefore, we numerically generated the normalized tpm values as described in this supplement and aggregated those values by dose. These are the exact same normalized tpm values that are aggregated by plate and dose and histogrammed in Fig. S3(g-i). Finally, for each dose we computed the 5% and 95% quantiles of the numerically generated normalized tpm values and plotted the results in Fig. 10(a,b,c) of the main text.

#### IV. ADDITIONAL DOSE RESPONSE AND CREDIBILITY REGION PLOTS.

In this section we provide additional dose-response and credibility region plots from analysis of experimental data, expanding on the three select genes shown in Fig. 10 of the main text.

We plot data for all 203 genes with small P-value for knockdown at dose=5, $P_{\mathrm{KD}_5} < 0.01$ in Figs. S4-S7. In order to format the data for plotting, it is sorted by MAP estimate of knockdown at dose=5 and is split into two batches consisting of 104 genes with lowest $\mathrm{KD}_5$'s (Figs. S4 & S6) and 99 genes with the higher$\mathrm{KD}_5$'s (Figs. S5 & S7). The gene depicted in Fig. 10(a,d,e) of the main text is gene 6827, it appears on row 7, column 2 of Figs. S4 & S6.

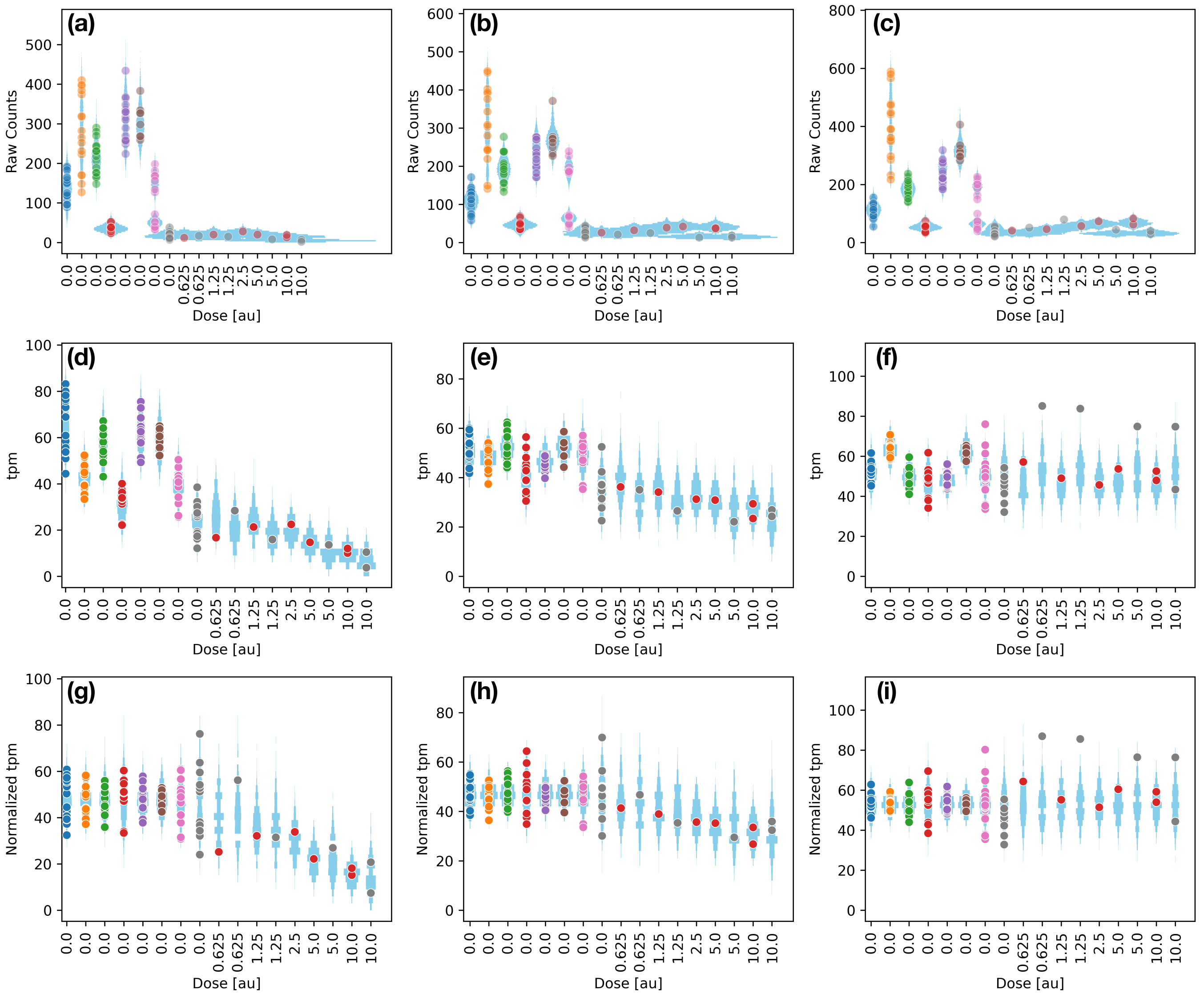


FIG. S3. Visualizing the MAP estimate of the negative binomial dispersion. The three columns of this plot correspond to the three columns of Fig. 10 of the main text and are plotted using the same data. The columns represent a strongly dose-responsive gene, a weakly dose-responsive gene, and a dose non-responsive gene. The rows represent different ways of visualizing gene expression data: raw counts (top row), tpm: transcripts per million (middle row), and tpm normalized to the fitted mean over the plates (bottom row). The colored dots represent experimental measurements, with the dot color indicating the plate index. Blue regions represent negative binomial distributions from MAP estimates (see Supplement text for details of what is plotted).

There are 3573 genes with intermediate P-values, $0.01 < P_{\mathrm{KD}_5} < 0.99$. We plot a random selection of 104 of these genes in Figs. S8 & S9. There are 4229 genes with large P-values, $0.99 < P_{\mathrm{KD}_5}$. We plot a random selection of 104 of these genes in Figs. S10 & S11.

We observe that the credibility regions for genes with low and intermediate P-values are consistently stretched along a line where $\mathrm{rIC}_{50}$ and mKD are anticorrelated. On the other hand credibility regions for genes with high P-values are clustered at mKD $\sim 1$, with little information on $\mathrm{rIC}_{50}$. We also observe that credibility regions for genes with lower P-values (and $\mathrm{KD}_5$'s) are more tight than those for genes with intermediate P-values.

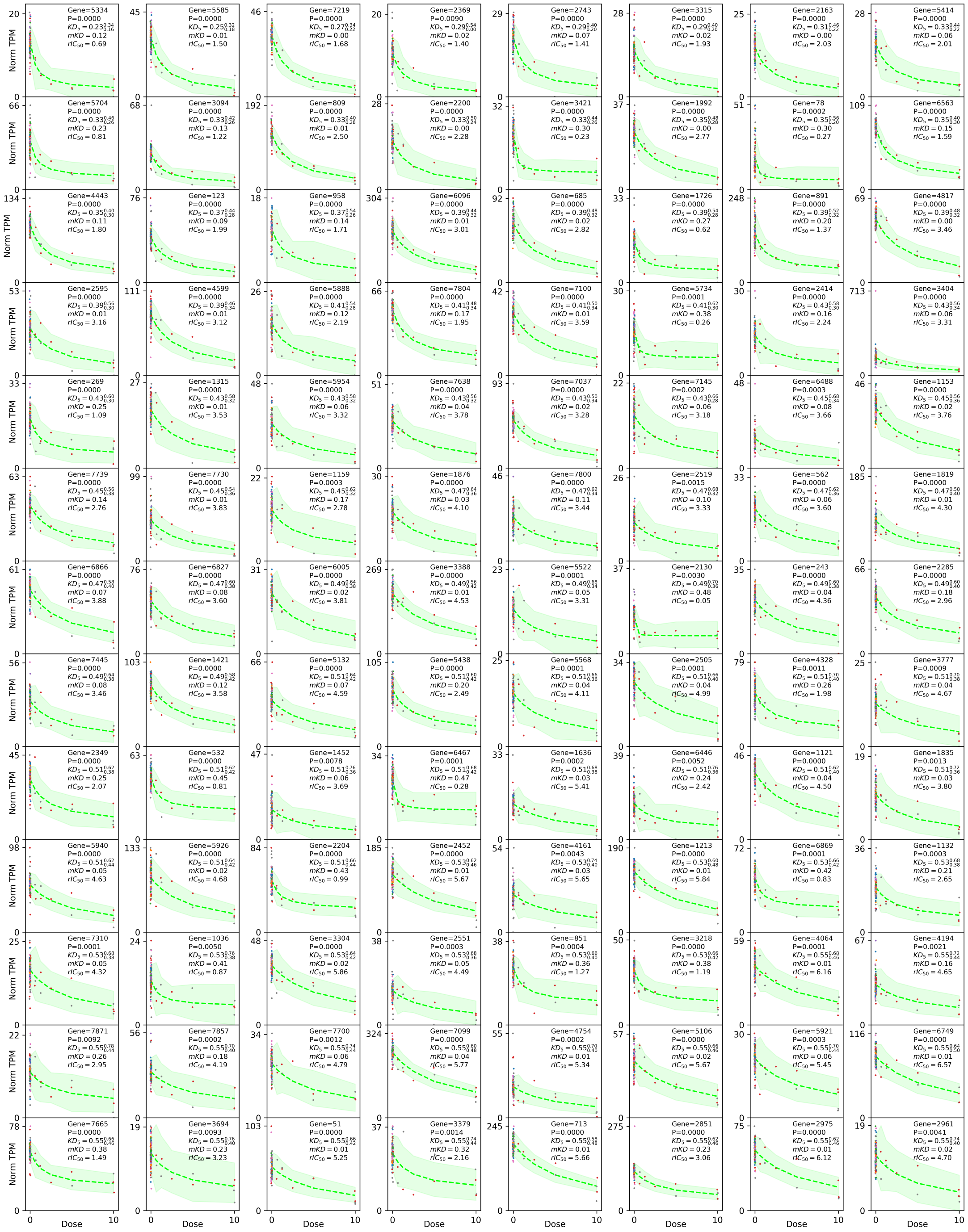

FIG. S4. First batch of dose-response fits for genes with small P-value for knockdown at dose=5, $P_{\mathrm{KD}_5} < 0.01$.

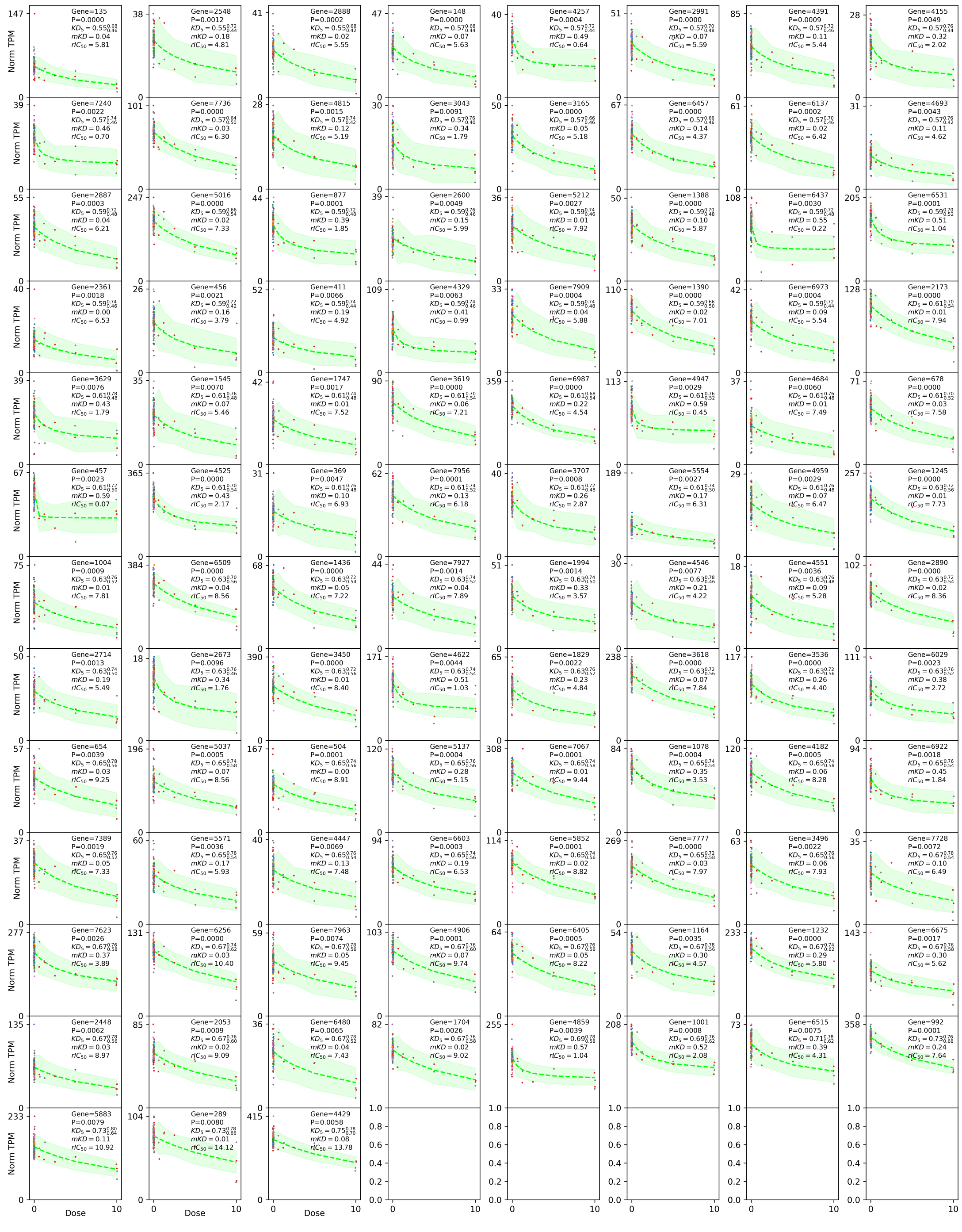

FIG. S5. Second batch of dose-response fits for genes with small P-value for knockdown at dose=5, $P_{\mathrm{KD}_5} < 0.01$.

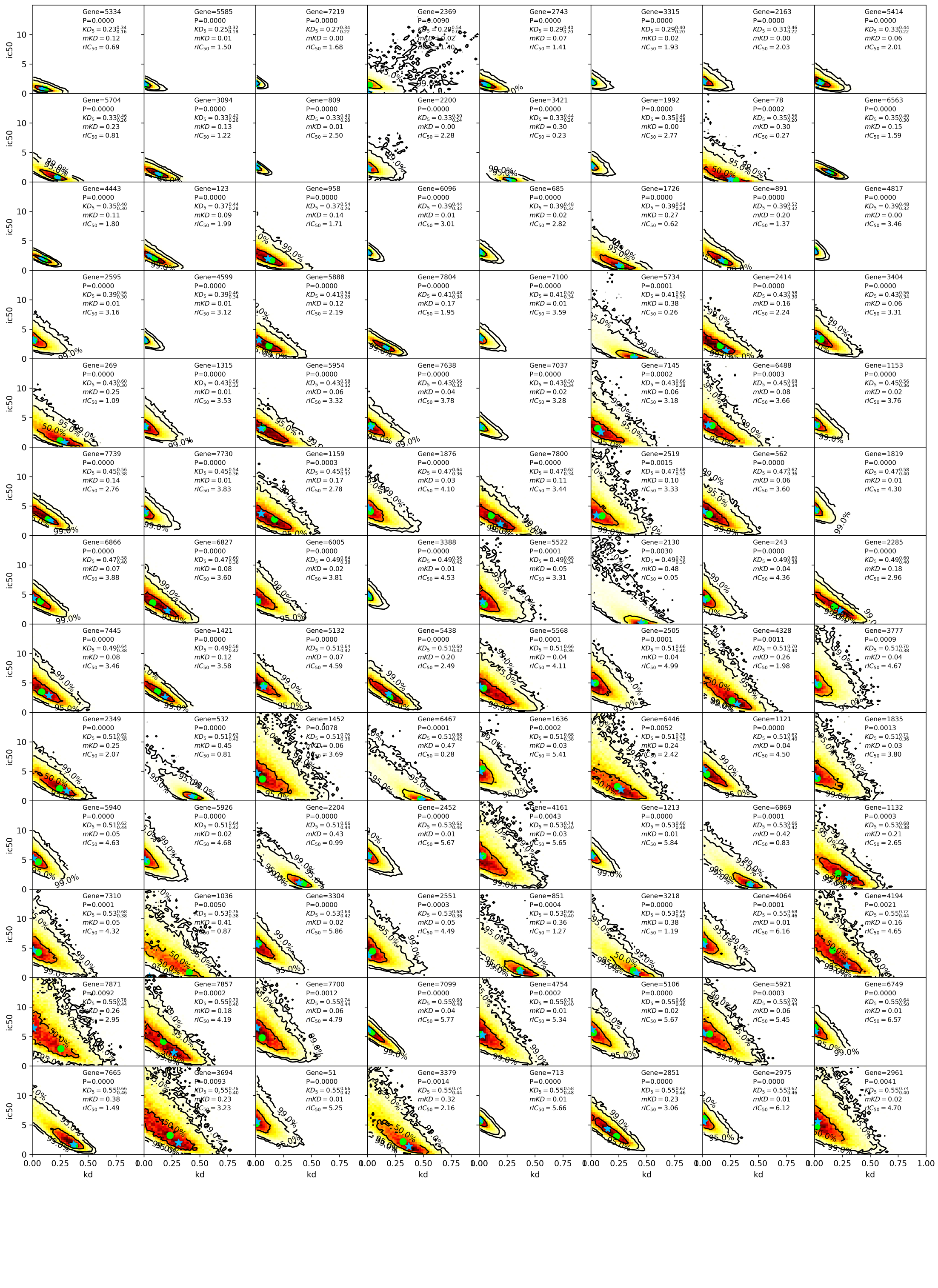

FIG. S6. First batch of credibility region plots for genes with small P-value for knockdown at dose=5, $P_{\mathrm{KD}_5} < 0.01$.

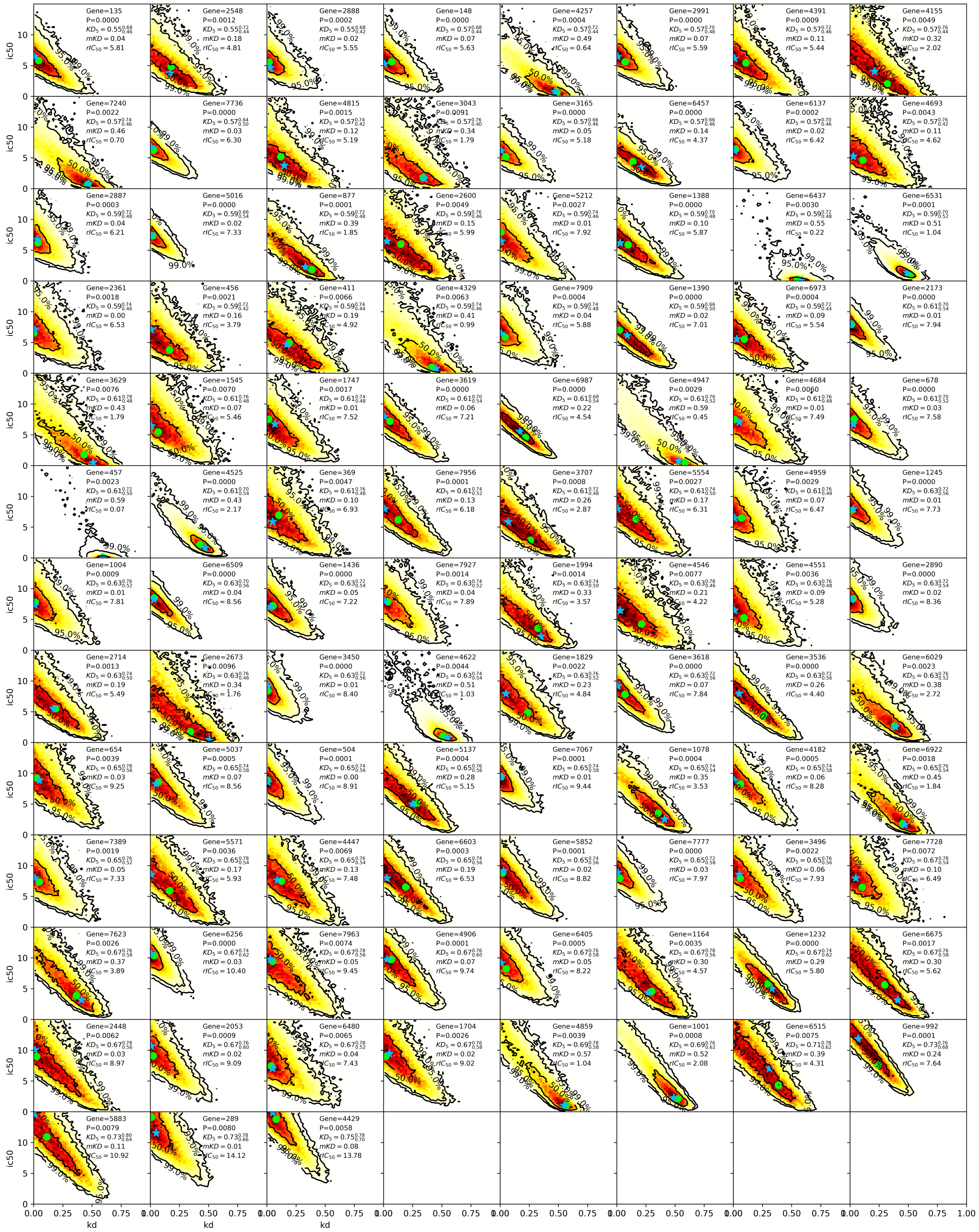

FIG. S7. Second batch of credibility region plots for genes with small P-value for knockdown at dose=5, $P_{\mathrm{KD}_5} < 0.01$.

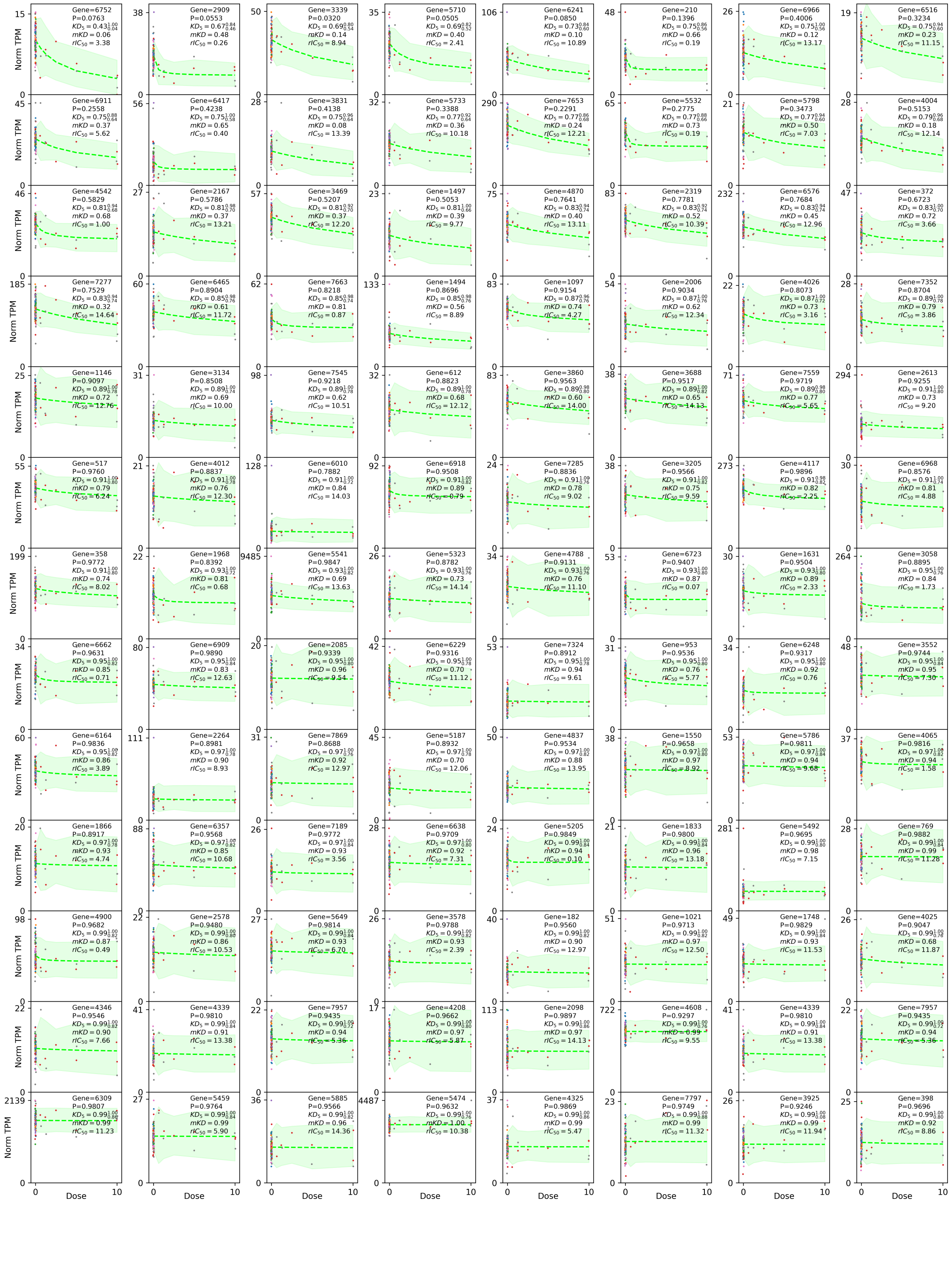

FIG. S8. Dose-response fits for genes with intermediate P-value for knockdown at dose=5, $0.01 < P_{\mathrm{KD}_5} < 0.99$.

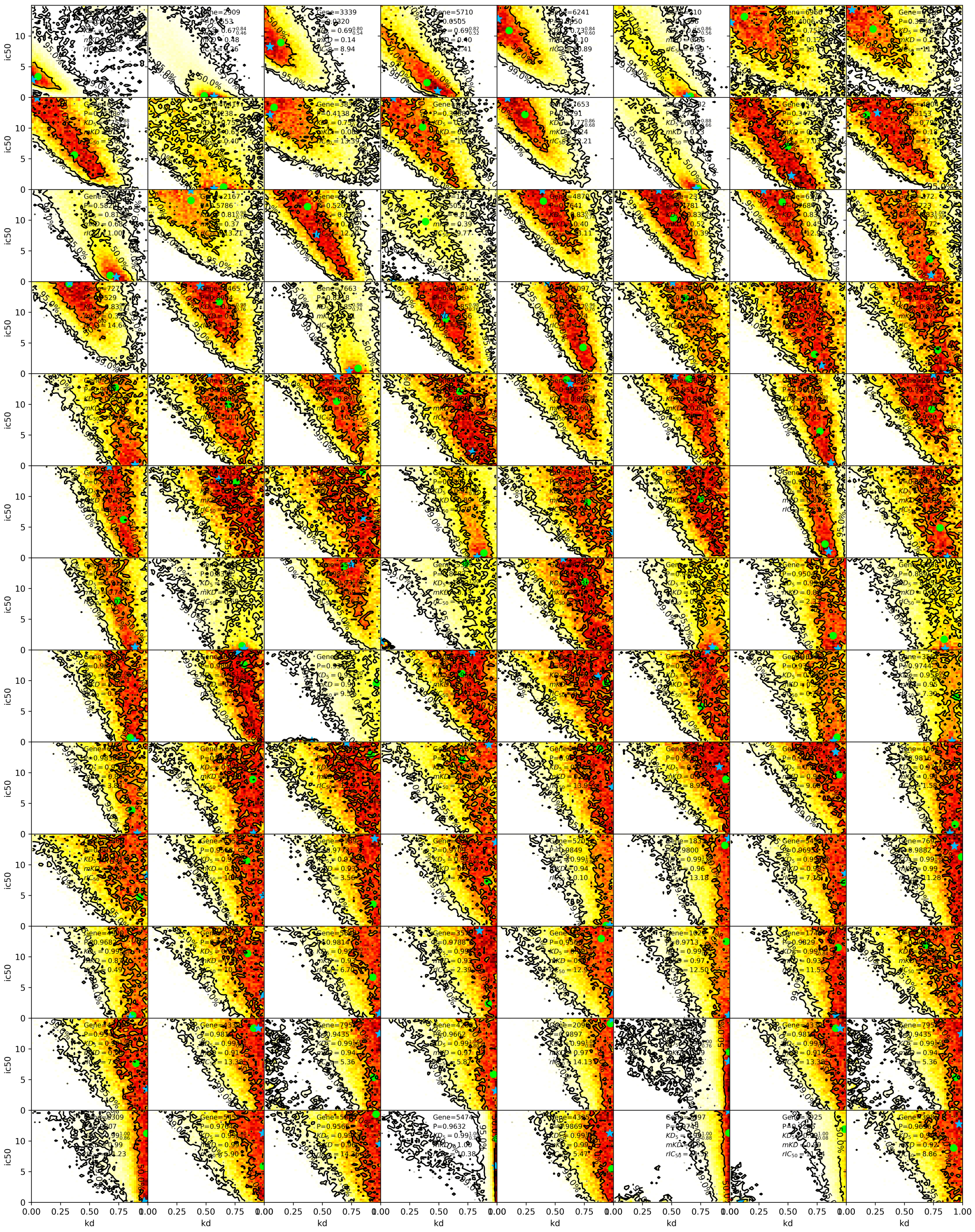


FIG. S9. Credibility region plots for genes with intermediate P-value for knockdown at dose=5, $0.01 < P_{\mathrm{KD}_5} < 0.99$.

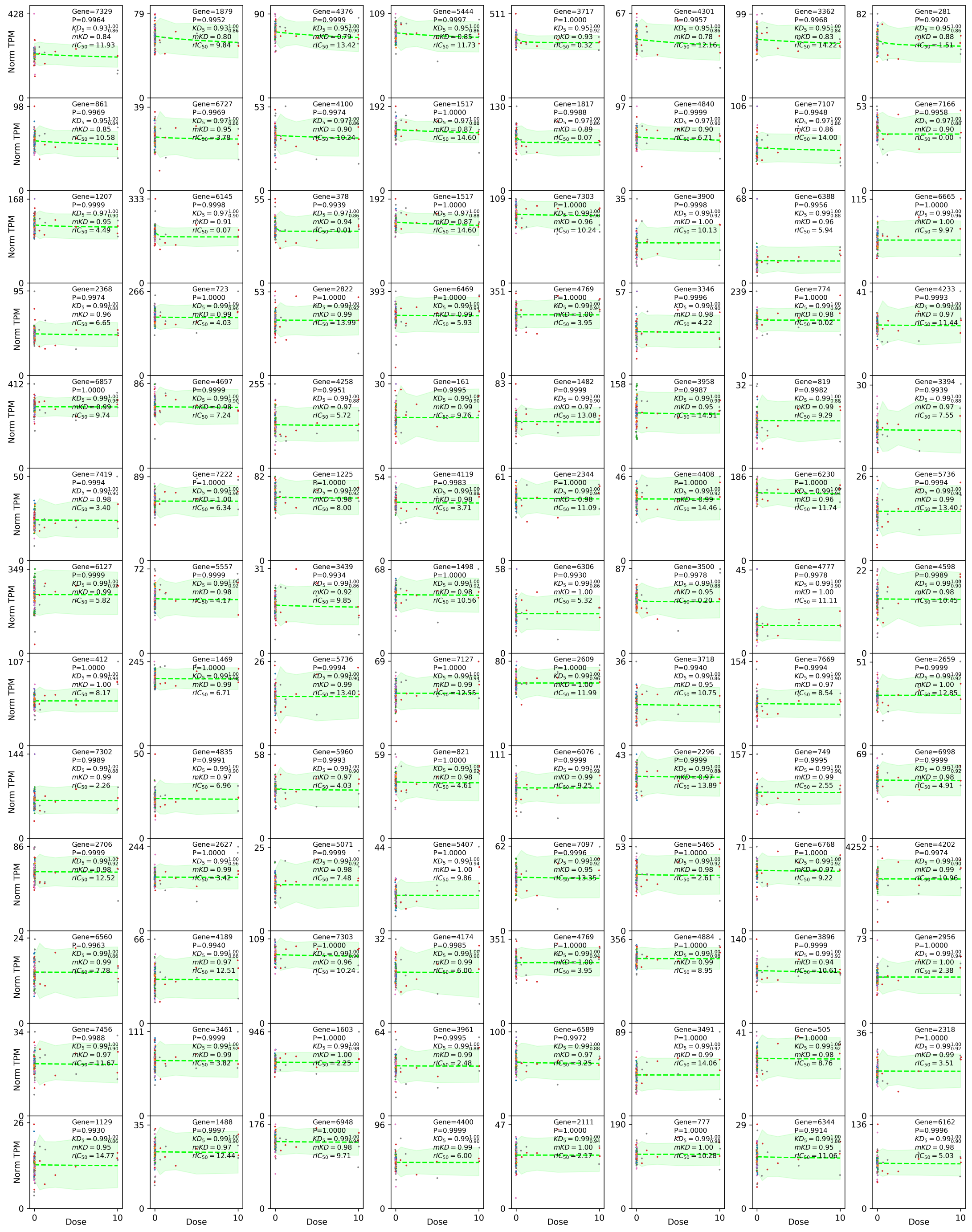

FIG. S10. Dose-response fits for genes with large P-value for knockdown at dose=5, $0.99 < P_{\mathrm{KD}_5}$.

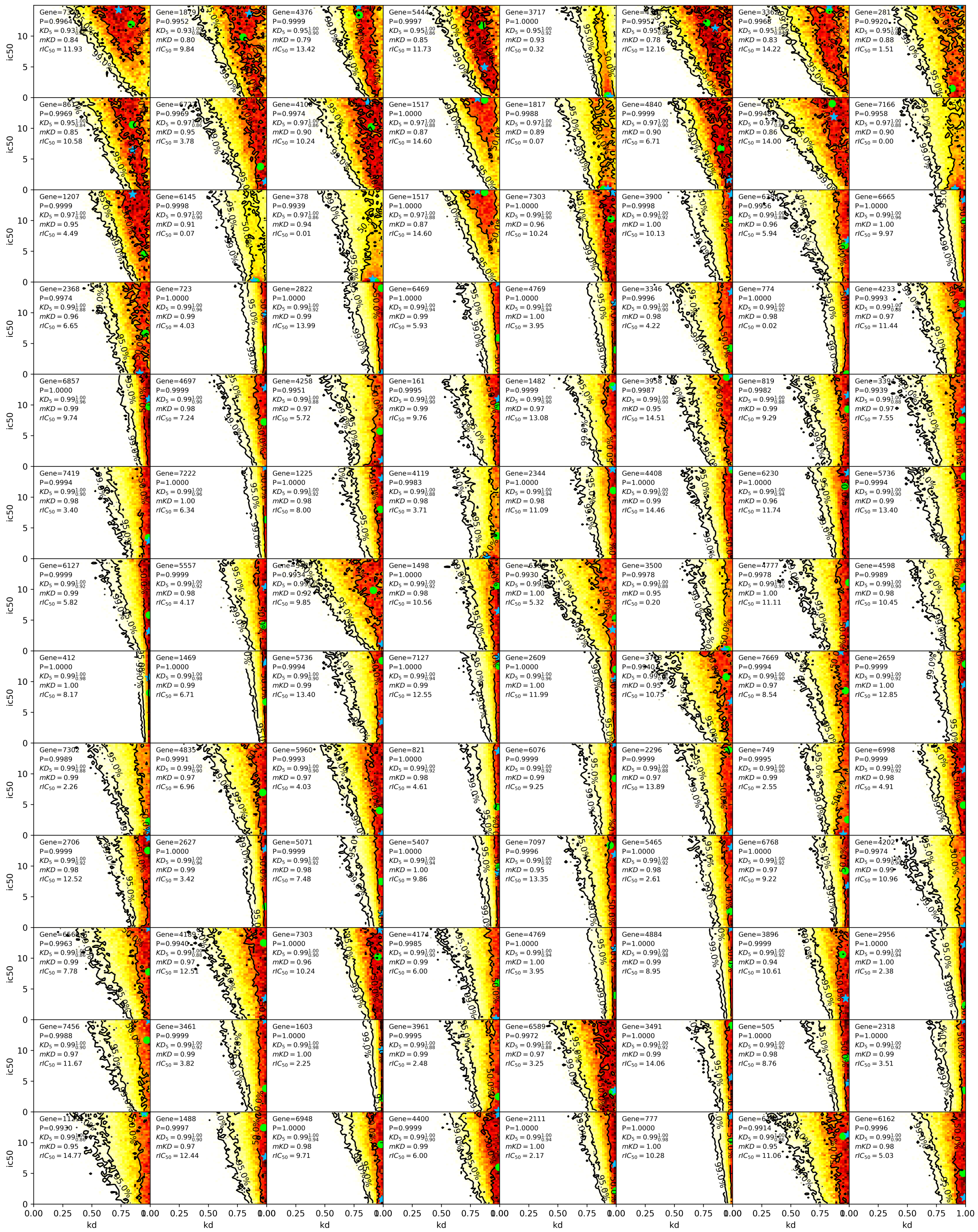

FIG. S11. Credibility region plots for genes with large P-value for knockdown at dose=5, $0.99 < P_{\mathrm{KD}_5}$.
